# Dataset complexity impacts both MOTU delimitation and biodiversity estimates in eukaryotic 18S rRNA metabarcoding studies

**DOI:** 10.1101/2021.06.16.448699

**Authors:** Alejandro De Santiago, Tiago José Pereira, Sarah L. Mincks, Holly M. Bik

**Affiliations:** Department of Marine Sciences, University of Georgia, 325 Sanford Drive, Athens, GA 30602, USA; Institute of Bioinformatics, University of Georgia, 120 Green Street, Athens, GA 30602, USA; School of Fisheries and Ocean Sciences, University of Alaska, Fairbanks, AK 99775, USA

**Keywords:** metabarcoding, OTUs, ASVs, 18S rRNA gene, microbial eukaryotes, nematodes, biodiversity

## Abstract

How does the evolution of bioinformatics tools impact the biological interpretation of high-throughput sequencing datasets? For eukaryotic metabarcoding studies, in particular, researchers often rely on tools originally developed for the analysis of 16S ribosomal RNA (rRNA) datasets. Such tools do not adequately account for the complexity of eukaryotic genomes, the ubiquity of intragenomic variation in eukaryotic metabarcoding loci, or the differential evolutionary rates observed across eukaryotic genes and taxa. Recently, metabarcoding workflows have shifted away from the use of Operational Taxonomic Units (OTUs) towards delimitation of Amplicon Sequence Variants (ASVs). We assessed how the choice of bioinformatics algorithm impacts the downstream biological conclusions that are drawn from eukaryotic 18S rRNA metabarcoding studies. We focused on four workflows including UCLUST and VSearch algorithms for OTU clustering, and DADA2 and Deblur algorithms for ASV delimitation. We used two 18S rRNA datasets to further evaluate whether dataset complexity had a major impact on the statistical trends and ecological metrics: a “high complexity” (HC) environmental dataset generated from community DNA in Arctic marine sediments, and a “low complexity” (LC) dataset representing individually-barcoded nematodes. Our results indicate that ASV algorithms produce more biologically realistic metabarcoding outputs, with DADA2 being the most consistent and accurate pipeline regardless of dataset complexity. In contrast, OTU clustering algorithms inflate the metabarcoding-derived estimates of biodiversity, consistently returning a high proportion of “rare” Molecular Operational Taxonomic Units (MOTUs) that appear to represent computational artifacts and sequencing errors. However, species-specific MOTUs with high relative abundance are often recovered regardless of the bioinformatics approach. We also found high concordance across pipelines for downstream ecological analysis based on beta-diversity and alpha-diversity comparisons that utilize taxonomic assignment information. Analyses of LC datasets and rare MOTUs are especially sensitive to the choice of algorithms and better software tools may be needed to address these scenarios.

## Introduction

Over the last ten years, there has been a rapid expansion of metabarcoding methods applied toward the study of eukaryotic taxa. These studies typically utilize loci such as the Cytochrome c Oxidase subunit I gene (COI) or other mitochondrial genes (e.g., 12S) for large vertebrates and macroinvertebrates (Arribas et al., 2016; Elbrecht & Leese, 2017; Hebert et al., 2003; Leray & Knowlton, 2016; Machida et al., 2012), the 18S or 28S ribosomal RNA (rRNA) nuclear genes for microbial invertebrates and single-celled eukaryotes (Creer et al., 2010; Pawlowski et al., 2012), ITS rRNA loci for fungi (Lindner et al., 2013; Toju & Baba, 2018), or the *rbcL* and *matK* plastid genes for plants and algae (Akita et al., 2019; Bell et al., 2017; Group et al., 2009; Wolf & Vis, 2020). Since metabarcoding studies utilize high-throughput sequencing (HTS) technologies (e.g., Illumina sequencing) and PCR primers that amplify a broad range of taxa from numerous samples at once, the resulting datasets have provided unprecedented biodiversity insights that have quickly expanded our understanding of spatio-temporal patterns and ecological interactions in diverse environments (Bik et al., 2012; Deiner et al., 2017; Zinger et al., 2019).

The massive amount of data produced by HTS platforms, however, requires stringent quality control steps to differentiate real biological reads from erroneous sequences. Proper study design is paramount for drawing robust biological and ecological conclusions from metabarcoding datasets (Alberdi et al., 2018; Murray et al., 2015). However, even the most well-designed studies can contain artefacts stemming from PCR, sequencing errors (e.g., Illumina “tag bleeding”), run batch effects (Fidler et al., 2020; Leek et al., 2012; Schnell et al., 2015), and microbial contamination (e.g., introduced via kit reagents, (Salter et al., 2014)). While some of these artefacts can be detected and bioinformatically eliminated via the incorporation of rigorous control samples (e.g., DNA extraction blanks, PCR negative controls, and mock community standards, (Alberdi et al., 2018; Deiner et al., 2017; Hornung et al., 2019)) and data cleaning software (e.g., Decontam, microDecon, (Davis et al., 2018; McKnight et al., 2019)), other “artefacts” appear to result from the biological nuances of the chosen metabarcoding locus (i.e., differential patterns of gene evolution across taxa). For example, intragenomic variation persists within nuclear rRNA loci, even though rRNA tandem repeat arrays are subjected to concerted evolution within eukaryotic genomes (Bik et al., 2013; Kumar et al., 2017; Pawłowska et al., 2020; Zhao et al., 2019). As a result, an individual eukaryote will typically be represented by a “Head-Tail” pattern in 18S rRNA metabarcoding datasets, exhibiting a dominant “Head” sequence with high relative abundance that represents the species-specific DNA barcode, as well as a “Tail” of rarer low-abundance sequences that exhibit high pairwise similarity to the diagnostic reference barcode (Porazinska, Giblin-Davis, Esquivel, et al., 2010; Porazinska, Giblin-Davis, Sung, et al., 2010).

The clustering of raw metabarcoding reads into Molecular Operational Taxonomic Units (MOTUs) is one ubiquitous approach for minimizing artefacts and reducing confounding biological variation such as intragenomic rRNA variation. MOTU is a general term for a DNA-based “species approximation” (Blaxter, 2016; Blaxter et al., 2005; Fonseca et al., 2012) that can be used to calculate traditional biodiversity indices and perform ecological analyses, either using taxa presence/absence or relative abundance. Two distinct classes of MOTUs seen in the scientific literature are associated with specific bioinformatics algorithms for processing raw HTS data: Operational Taxonomic Units (OTUs) and Amplicon Sequence Variants (ASVs). OTUs emerged as an early approach for analyzing data from diverse HTS platforms, relying on the distance-based clustering of raw reads according to a set pairwise similarity cutoff (e.g., 97% and 99% for prokaryotes and eukaryotes, respectively (Deiner et al., 2017)). A plethora of OTU clustering algorithms now exist which can cluster reads in myriad ways (Jackson et al., 2016; Prodan et al., 2020), with endless parameter choices for the strictness (or inclusivity) of the OTU cluster generation steps. Two highly popular OTU clustering algorithms include UCLUST (Edgar, 2010) and VSearch (Rognes et al., 2016), respectively implemented in the QIIME1 (Caporaso et al., 2010) and QIIME2 (Bolyen et al., 2019) software suites for microbial ecology. Newer ASV algorithms utilize Illumina error correction approaches to perform quality checks and processing on raw sequence reads. Currently, the two most popular tools for ASV generation are DADA2 (Callahan et al., 2016) and Deblur (Amir et al., 2017), although other ASV algorithms are rapidly emerging (e.g., UNOISE2, (Edgar, 2016)). The most important distinction between these two classes of MOTUs is the increased sophistication of ASV algorithms, which include complex mathematical modelling approaches aimed at eliminating false positives, chimeras, and sequencing artefacts (although note that more simplistic chimera checking steps are also ingrained in many OTU clustering pipelines, (Edgar et al., 2011; Mysara et al., 2017)). Furthermore, ASVs are thought to represent the true DNA sequences present in the target community, and are thus reusable and reproducible across metabarcoding studies (in contrast to *de novo* OTUs which are emergent features of a dataset and must be treated as study-specific clusters, (Callahan et al., 2017)).

The impact of different MOTU generation workflows, and the biological implications for the move towards ASVs, has not been extensively documented for eukaryotes. The vast majority of benchmarking studies and algorithm comparisons have been performed on 16S rRNA marker gene studies of bacterial and archaeal communities (Caruso et al., 2019; Jackson et al., 2016; Nearing et al., 2018; Prodan et al., 2020). Although these prokaryotic-focused studies have provided important insights on the performance and ecological relevance of different computational workflows, their relevance for eukaryotic taxa is not always clear. Eukaryotic genome complexity and evolution may pose challenges for bioinformatics algorithms originally optimized for use with bacterial/archaeal 16S rRNA datasets. Additionally, many eukaryotic metabarcoding studies include complementary visual surveys of taxa (e.g., microscopy, trawls, kick sampling, video transects), which can provide critical independent observations for validating DNA-based inferences derived from HTS datasets (Cahill et al., 2018; Dell’Anno et al., 2015; Djurhuus et al., 2018; Geisen et al., 2018; Schuelke et al., 2018).

The current eukaryotic metabarcoding literature is heavily focused on methods comparisons for field sampling (Beentjes et al., 2019; Koziol et al., 2019; Turner et al., 2015) and wet lab protocols (e.g., DNA extraction comparisons, design of improved primer sets, (Alberdi et al., 2018; Bradley et al., 2016; Brannock & Halanych, 2015)), as well as comparison of intraspecific vs. interspecific genetic diversity for the COI gene (Elbrecht et al., 2018; Leray et al., 2013; Macheriotou et al., 2019). Yet, methods comparisons of eukaryotic metabarcoding workflows and benchmarking of other genetic loci (i.e., 18S rRNA and ITS rRNA datasets) are still limited, particularly when assessing the performance of algorithms producing distinct classes of MOTUs (i.e., OTUs vs. ASVs, (Bálint et al., 2014; Macheriotou et al., 2019; Pauvert et al., 2019)). Thus, there is a pressing need for downstream comparisons of bioinformatics tools in eukaryotic metabarcoding studies, especially those that evaluate the biological implications of different MOTU generation approaches and how the choice of software tools impacts downstream ecological analyses.

In the present study, we aimed to evaluate how distinct bioinformatics pipelines may impact the biological inferences drawn from 18S rRNA metabarcoding datasets. We focused on four computational workflows representing two distinct classes of MOTU generation (Figure 1): OTU clustering using VSearch and UCLUST, and ASV generation using DADA2 and Deblur. Our study compared results from two 18S rRNA metabarcoding datasets generated as part of other ongoing projects: a “high complexity” (HC) environmental dataset generated from bulk community DNA in Arctic marine sediments, and a “low complexity” (LC) dataset representing metabarcoding profiles from individually-barcoded nematodes, where morphological identification of each specimen was verified under light microscopy (Schuelke et al., 2018). Both datasets represent the same 18S rRNA gene region (V1–V2 hypervariable regions) and were generated using the same PCR primer set and molecular wet lab protocols, thus facilitating direct comparison of downstream bioinformatics outputs. We hypothesized that 1) ASVs would represent a more biologically relevant approach for eukaryotic metabarcoding data, 2) Species-specific DNA barcodes (Head MOTUs exhibiting high relative abundance) would be consistently recovered across all bioinformatics pipelines, 3) Computational algorithms and parameters would strongly influence downstream estimates of alpha- and beta-diversity, and 4) Dataset complexity (i.e., the level of biodiversity contained in each metabarcoding sample) would impact the performance of bioinformatics pipelines in unanticipated ways.

**Figure 1.**
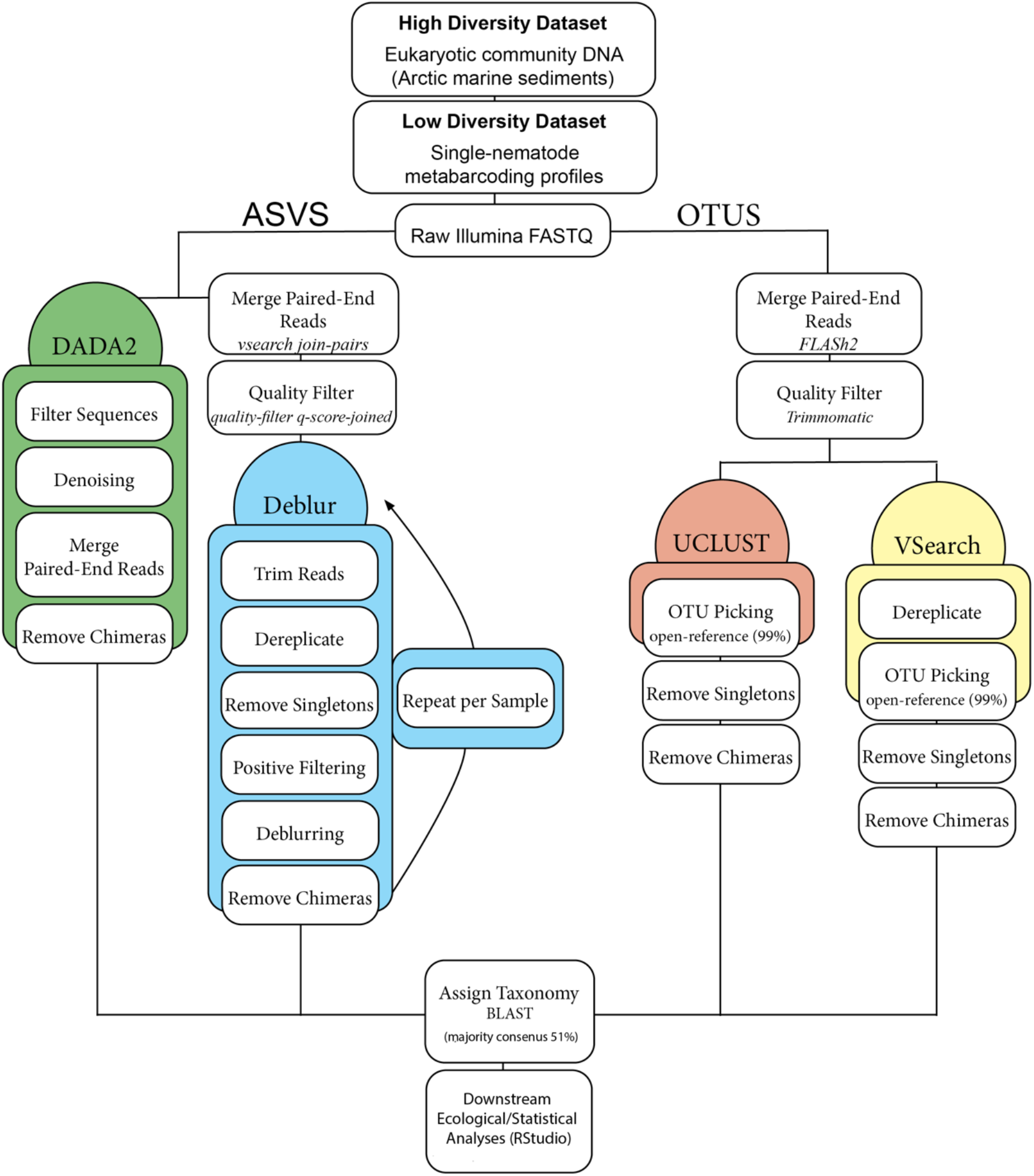
Workflow diagram of the four different bioinformatics pipelines used in this study. For the UCLUST and VSearch pipelines, reads were first quality controlled and merged using FLASH2 and Trimmomatic. Reads were clustered into OTUs using a 99% similarity threshold. Singletons and chimeric sequences were removed. For DADA2, the demultiplexed FASTQ files were used to estimate ASVs. For Deblur, denoising was done on merged quality-filtered reads. Unlike the DADA2 algorithm, Deblur processes each sample independently. Four matrices were produced for each dataset. Taxonomy was assigned using BLAST+ (BLAST majority consensus) in QIIME2 v2019.4.

## Materials and Methods

### Sample Collection and Data Generation

The high complexity (HC) Arctic metabarcoding dataset was generated using 127 marine sediment samples collected from the continental shelf/slope in the Northeast Chukchi and Beaufort Seas (Figure S1, Table 1). This dataset was generated as part of a broad study focused on evaluating benthic meiofauna community structure through a combination of morphological taxonomy and -Omics approaches. Additional details on sediment sample collection and sample processing are provided in Mincks et al. (2021). Briefly, frozen sediment samples were thawed and meiofaunal organisms were isolated via decantation over a 63 μm mesh sieve. Total genomic DNA was extracted from material retained on the sieve using MoBio PowerSoil® kits (MoBio Laboratories, Inc., Carlsbad, CA). A fragment of the 18S rRNA gene (∼400-bp, V1–V2 hypervariable regions) was amplified from environmental DNA extracts using the F04/R22 eukaryotic primers (Creer et al., 2010). PCR products from each sediment sample were tagged with a unique nucleotide barcode and pooled before sequencing.

**Table 1.** Description of geographic locations and number of samples included in this study.

| **Alaskan Continental Shelf**<br>**(High Complexity Dataset)** | **Number of**<br>**Samples** | **Region** | **Depth Range (m)** |
| --- | --- | --- | --- |
| Alaskan Beaufort Shelf | 8 | Arctic | 50-500 |
| Amundsen Gulf | 30 | Arctic | 32-352 |
| Banks Island | 7 | Arctic | 51-379 |
| Camden Bay | 13 | Arctic | 20-350 |
| Chukchi Sea | 32 | Arctic | 25.4-51.1 |
| Mackenzie River Plume | 37 | Arctic | 17-1200 |
| **Nematode Microbiomes**<br>**(Low Complexity Dataset)** | **Number of**<br>**Samples** | **Region** | **Depth Range (m)** |
| Chromadoridae | 31 | Arctic, Gulf of Mexico,<br>Southern California | 0-1000 |
| Comesomatidae | 46 | Arctic, Gulf of Mexico | 20 -1000 |
| Desmoscolecidae | 29 | Arctic, Gulf of Mexico | 200 - 1000 |
| Oxystominidae | 27 | Arctic, Gulf of Mexico | 20 - 1000 |
| Other | 94 | Arctic, Gulf of Mexico,<br>Southern California | 0 - 2239 |

The low complexity (LC) single-specimen metabarcoding dataset was generated from individual marine nematodes collected in the Pacific, Arctic, and Gulf of Mexico (Figure S1, Table 1). This dataset was originally generated by Schuelke et al. (2018) who evaluated microbiome patterns (archaea and bacteria) associated with diverse marine nematode genera. Further exploration of intragenomic rRNA patterns in marine nematodes was carried out by Pereira et al. (2020) in a refined version of the original dataset (i.e., 227 samples). Both of these studies provide detailed methods regarding nematode sorting, taxonomic identification, and genus-level diversity patterns. Wet lab protocols for single-nematode metabarcoding included an extensive suite of blank/control samples as a checkpoint for contamination in these low biomass samples. After morphological vouchering of each nematode specimen, samples were submitted to molecular procedures described in Pereira et al. (2020). Briefly, DNA was extracted from individual nematodes using a Proteinase K “Worm Lysis Buffer” protocol. PCR amplification of the 18S rRNA gene was carried out using the same eukaryotic primer set (F04/R22; Creer et al., 2010) and multiplexing/pooling procedures as described for the HC dataset above.

All PCR products were purified using magnetic beads following the manufacturer’s protocol (Agencourt AMPure XP beads; Beckman Coulter). Sample concentrations were subsequently measured using a Qubit® 3.0 Fluorometer and a Qubit® dsDNA HS (High Sensitivity) Assay Kit (Thermo Fisher Scientific). Normalization values were calculated to ensure that approximately equivalent DNA concentrations were pooled across all samples, including controls and blank samples. The final pooled libraries were subjected to an additional magnetic bead cleanup step, followed by size selection on a BluePippin (Sage Science) to remove any remaining primer dimer and isolate target PCR amplicons within the range of 300–700 bp. A Bioanalyzer trace was run on each size-selected pool as a quality control measure, and the pooled 18S rRNA amplicon libraries were sequenced in two separate runs on the Illumina MiSeq platform (2 × 300-bp paired-end runs) at the UC Davis Genomics Core Facility. All wet laboratory protocols, sample mapping files, and downstream bioinformatics scripts used in this study have been deposited on GitHub (https://github.com/BikLab/OTU-ASV-euk-benchmarking).

### OTU and ASV Pipeline Designs

OTU clustering was carried out using the UCLUST and VSearch algorithms, while ASVs were generated using the DADA2 and Deblur pipelines (Figure 1). For OTU picking workflows, raw Illumina reads were first demultiplexed, merged using FLASH2 (Magoč & Salzberg, 2011), and quality filtered using Trimmomatic (Bolger et al., 2014) in conjunction with a custom script described in Schuelke et al. (2018). UCLUST (Edgar, 2010) was implemented within QIIME1 v1.9.1 (Caporaso et al., 2010), whereby quality-filtered reads were clustered at 99% using the subsampled open-reference protocol (*pick_open_reference_otus.py*), with 10% subsampling of failed reads, and discarding of singletons from the final OTU table (Rideout et al., 2014). Chimeric sequences were identified and removed from the OTU table using USEARCH61, and the resulting OTUs were aligned to the QIIME-formatted SILVA132 database using the Pynast aligner and UCLUST (pairwise alignment method). Highly variable regions were removed using *filter_alignment.py*. Next, aligned representative sequences were used to construct a phylogeny using Fasttree (Price et al., 2009) and rooted using the midpoint method (*make_phylogeny.py*). VSearch (Rognes et al., 2016) was implemented using QIIME2 v2019.4 (Bolyen et al., 2018), whereby quality-filtered reads were clustered at 99% using *cluster-features-open-reference*. Singletons were filtered out from the OTU table and chimeric sequences identified using the *de novo* option; both chimeric and borderline chimeric sequences were removed from the OTU table. The resulting representative sequences were aligned using MAFFT (Katoh et al., 2002), ambiguous regions were masked, and the resulting aligned sequences were used to infer a phylogenetic tree using Fasttree (*align-to-tree-mafft-fasttree*, with rooting using the midpoint method). Discrepancies between UCLUST and VSearch workflows were a result of algorithm changes between the QIIME1 and QIIME2 pipelines (e.g., MAFFT replacing the depreciated Pynast aligner in QIIME1, and UCLUST being replaced with VSearch in QIIME2). The UCLUST and VSearch workflows thus aimed to carry out the same approximate bioinformatics steps in QIIME1 vs. QIIME2, respectively. It was therefore crucial to assess whether these pipeline changes in QIIME2 would result in noticeable discrepancies in downstream results and biological and ecological interpretations.

For ASV workflows, DADA2 (Callahan et al., 2016) and Deblur (Amir et al., 2017) were both executed within QIIME2 v2019.4. DADA2 was implemented directly on the demultiplexed FASTQ files since this algorithm incorporates quality-filtering of raw Illumina reads. The DADA2 error-correcting algorithm was run using default parameters, except for the truncating and trimming parameters. Forward and reverse reads were truncated at 220bp and 236bp for the HC dataset and at 232bp and 253bp for the LC dataset (median PHRED score ≥ 30). Unlike DADA2, Deblur does not differentiate between forward and reverse reads. Therefore, reads were initially merged using the *vsearch join-pairs* command and quality-filtered according to the *q-score-joined* protocol. Deblur was implemented on the merged quality-controlled reads using default parameters. Reads were trimmed at 360bp and 350bp for the HC and LC dataset, respectively, thus allowing us to maximize the initial number of reads for the Deblur pipeline. Deblur internally removes chimeric and error-prone reads on a sample-by-sample basis. A tree was generated for the DADA2 and Deblur datasets using the *align-to-tree-mafft-fasttree* workflow in QIIME2 as described above (see VSearch method).

### Bioinformatics Analysis

All downstream analyses were conducted in RStudio (Team, 2017) using the phyloseq (McMurdie & Holmes, 2013), vegan (Oksanen, 2011), ggplot2 (Wickham, 2009), and ggpubr (Kassambara, 2018) packages. First, we recorded the overall number of reads and OTUs/ASVs retained by each pipeline (Tables S1, S2), including comparisons between individual samples and datasets (HC and LC). In order to assess whether the number of MOTUs was correlated with the number of reads retained by each pipeline, we estimated correlations (R^2^ and p-value) using Pearson’s correlation coefficient (Figure 2).

**Figure 2.**
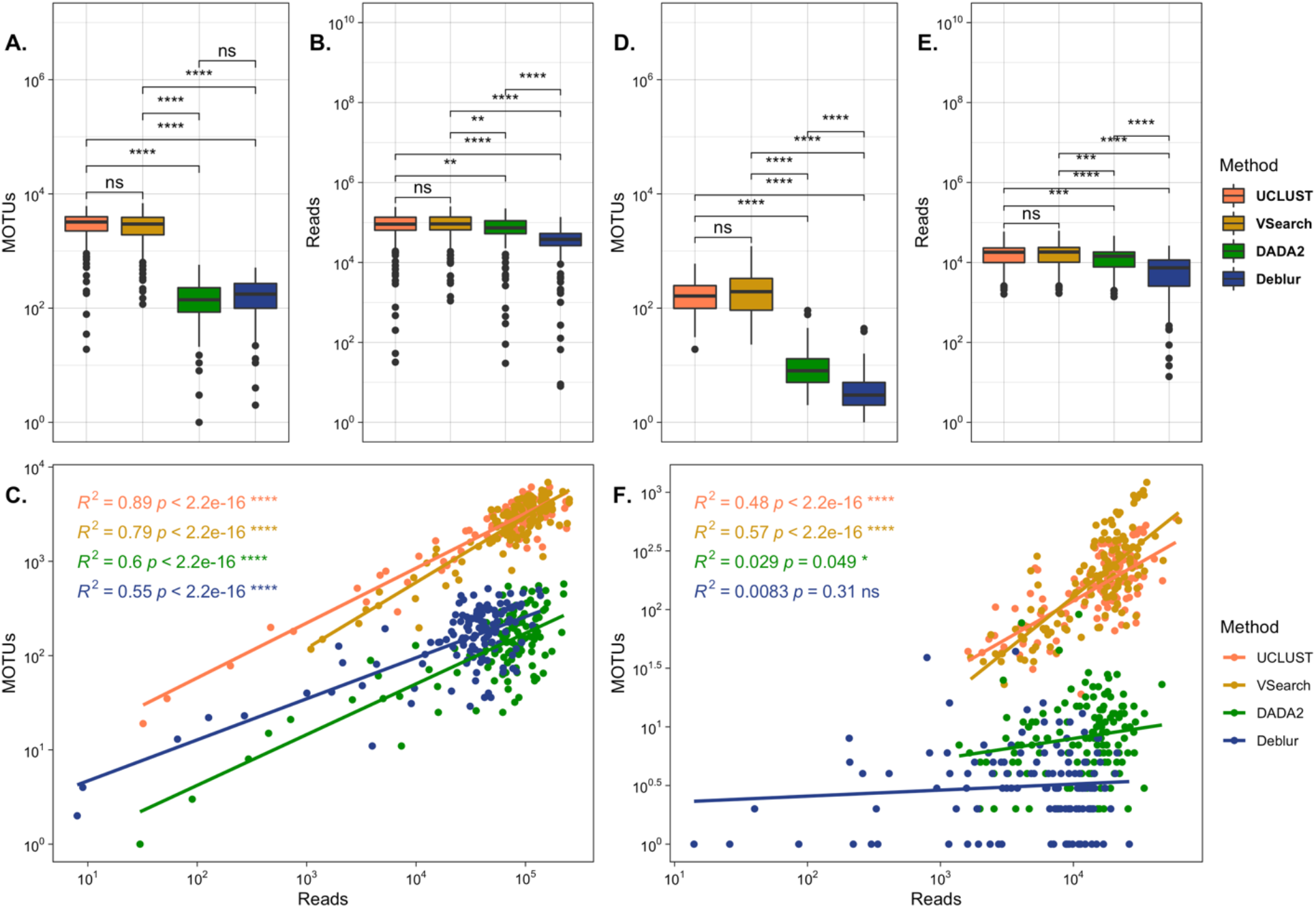
Number of MOTUs and reads retained by UCLUST, VSearch, DADA2, and Deblur. Median number of MOTUs (A, D) and reads (B, E) for HC and LC datasets, respectively. (C, F) Relationship between the number of MOTUs and number of reads across the four bioinformatics pipelines for HC and LC datasets, respectively.

Second, we aimed to evaluate whether each pipeline resulted in similar alpha- and beta-diversity metrics across Arctic subregions (HC dataset) and nematode taxonomic groups (LC dataset). Alpha-diversity was calculated in phyloseq using three different indices (observed MOTUs, Simpson, and Shannon) and visualized using ggplot2 and ggpubr (Figures S2, S3, S4). Kruskal-Wallis analysis was used to test for significant differences in alpha-diversity among Arctic subregions and nematode groups (i.e., most common nematode families). Pairwise comparisons were performed using the Wilcox test when significant differences among groups were detected. Principal Coordinate Analysis (PCoA) was carried out to assess beta-diversity using three different metrics: Bray-Curtis similarity, weighted Unifrac, and unweighted Unifrac (Lozupone & Knight, 2005). The HC dataset was rarefied at 1000 sequences per sample, resulting in five samples that were excluded from the UCLUST, DADA2, and Deblur pipelines (Table S1). The LC dataset was also rarefied at 1000 sequences per sample, resulting in 22 samples that were excluded from the Deblur pipeline (Table S2). Finally, a Procrustes analysis was used to determine concordance among the MOTU pipelines and beta-diversity metrics. Procrustes were visualized using a combination of vegan and ggplot2 packages. The Procrustes results were further assessed using PROTEST with 9999 permutations (Jackson, 1995). Procrustes and PROTEST were implemented on rarefied datasets (Table 2). For each comparison using Procrustes/PROTEST, samples not present in both pipelines were excluded from the analysis (i.e., samples that did not meet the minimum rarefaction threshold). The impact of bioinformatics parameters and distance indices was assessed using the returned R and M^12^ values from each pairwise comparison. Outputs are considered to be highly concordant if two algorithms displayed a high R value alongside a low M^12^ value, whereas the opposite (i.e., low R value alongside a high M^12^ value) characterizes them as being more discordant.

**Table 2.**
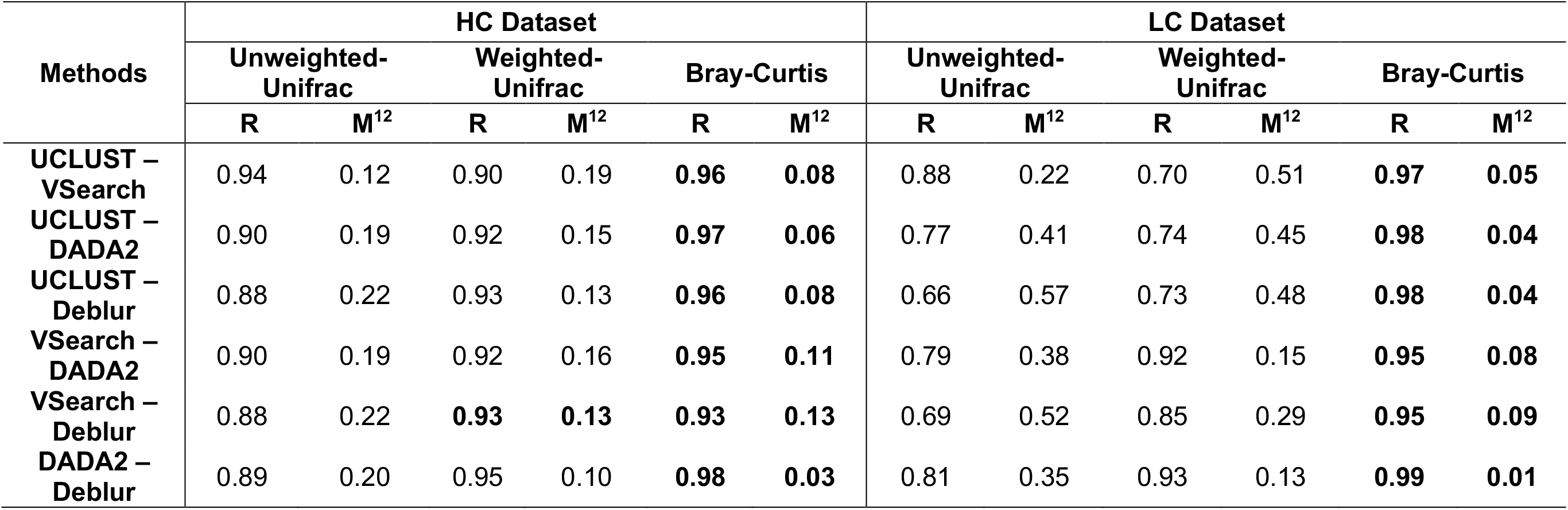
Results from the PROTEST/Procrustes analysis. For each dataset (i.e., distance index), the highest values of concordance (R: correlation coefficient and M^12^: goodness-of-fit statistic) are highlighted in bold. Beta diversity was estimated by rarefying data matrices at 1000 reads per sample. For all comparisons, p-value was always significant (*P* < 0.01).

Taxonomy was assigned for each MOTU table using QIIME2 v2019.4. For UCLUST, FASTA files containing OTU representative sequences were reformatted for compatibility with QIIME2. For all four pipelines, taxonomy was assigned using BLAST+ with a minimum of 90% sequence identity. Our reference database of full-length 18S rRNA sequences was the QIIME-formatted SILVA 132 database with additional curated 18S rRNA sequences, as described in Pereira et al. (2020). The impact of the MOTU pipelines on taxonomic assignment was further explored by generating barplots with ggplot2 of the ten most abundant taxonomic groups (order and family levels for HC and LC datasets, respectively; Figure 3).

**Figure 3.**
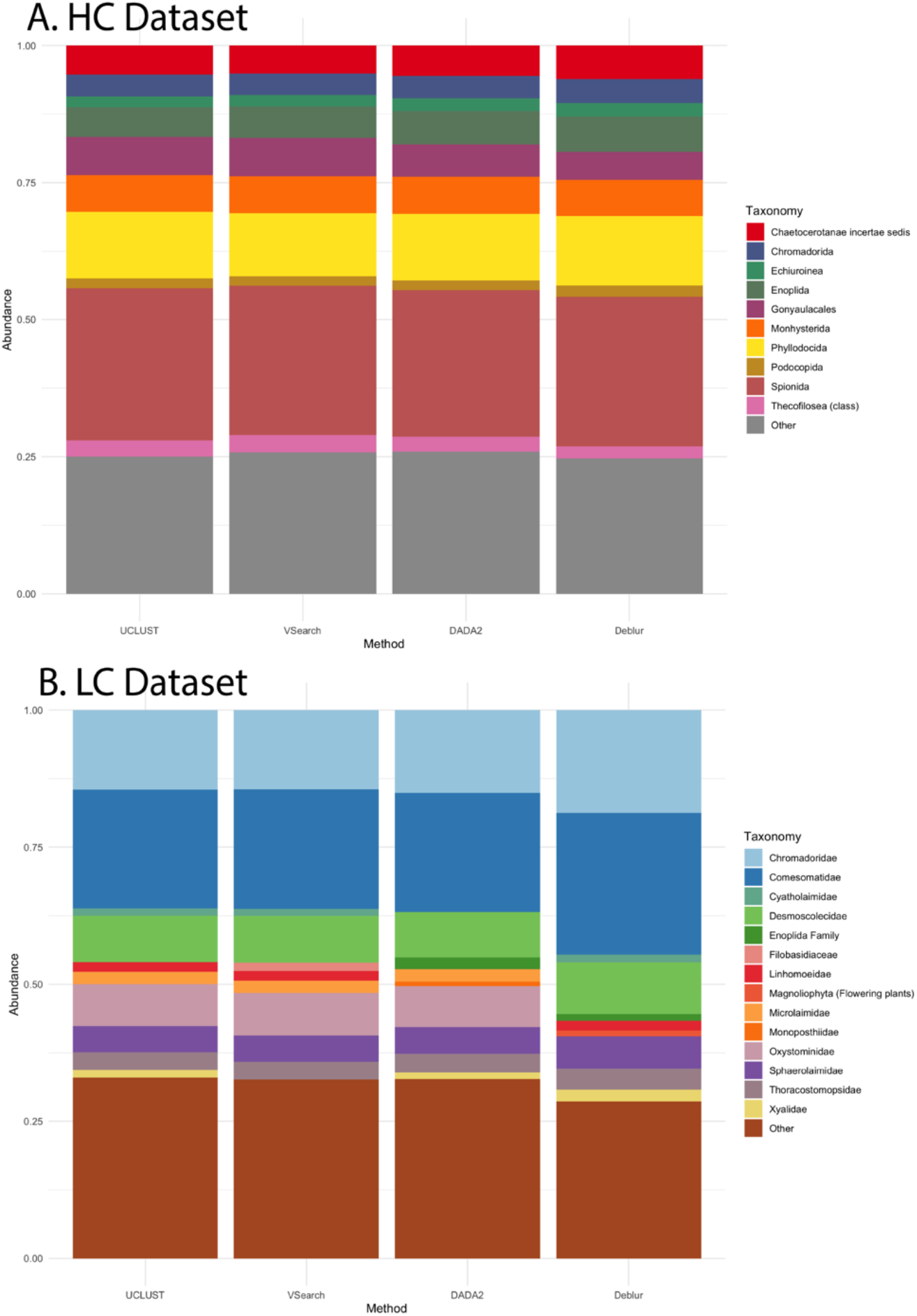
Barplots showing the top 10 most abundant taxa (i.e., relative abundance) across the different bioinformatics pipelines for (A) HC and (B) LC datasets. Taxonomic ranks are Order (HC-dataset) and Family (LC-dataset) levels. Lower abundance taxa were grouped in the category “Other”. Note that the top 10 most abundant taxa do not necessarily match across pipelines.

## Results

### Comparison of MOTUs recovered across bioinformatics pipelines

The total number of processed reads and MOTUs, including summary statistics (i.e., mean, median, standard deviation, and minimum/maximum values) obtained across the four pipelines for the HC and LC datasets are presented in Tables S1, S2, respectively. For both datasets, the number of processed reads and MOTUs varied across bioinformatics pipelines, especially when comparing between algorithm classes (i.e., ASVs vs. OTUs; Figure 2). For processed Illumina reads, significant differences were detected between pipelines (Global Kruskal-Wallis, < 0.01, LC X^2^ = 111.12; HC X^2^ = 110.29), except when comparing UCLUST vs. VSEARCH (not significant in either the HC or LC dataset; Figure 2B, 2E). Furthermore, while three pipelines resulted in a similar number of final processed reads (UCLUST, VSearch, and DADA2), the Deblur algorithm returned far fewer reads and recovered only about 43-57% of the processed reads seen in the other pipelines (Tables S1, S2).

The median number of MOTUs produced by OTU pipelines (UCLUST and VSearch) was at least fifteen-fold higher than the number of ASVs resulting from either DADA2 or Deblur (Figure 2A, 2D). For both datasets, there were no significant differences in the median number of MOTUs produced by UCLUST and VSearch. However, we observed subtle differences in OTU membership: VSearch consistently returned a higher proportion of “rare” OTUs (i.e., containing ≤ 10 reads per OTU; Table 3). This higher level of rare OTUs was much more pronounced for the LC dataset, with VSearch returning 39,967 more rare OTUs than UCLUST (a 266% increase; Table 3). For the HC dataset, VSearch exhibited 27,538 more rare OTUs than UCLUST (a 137% increase; Table 3). The MOTU patterns for ASV algorithms showed stark differences across the HC and LC datasets. For the Deblur pipeline, the median number of ASVs observed in the LC dataset was significantly lower than DADA2 (p-value = < 0.001, Figure 2D). In contrast, Deblur returned a higher median number of ASVs in the HC dataset, but not significantly different from DADA2 (Figure 2A). Although the number of Deblur ASVs appeared to be impacted by data complexity, the Deblur pipeline always returned lower total numbers of processed reads and MOTUs in each overall dataset compared to DADA2 (Table 3).

**Table 3.**
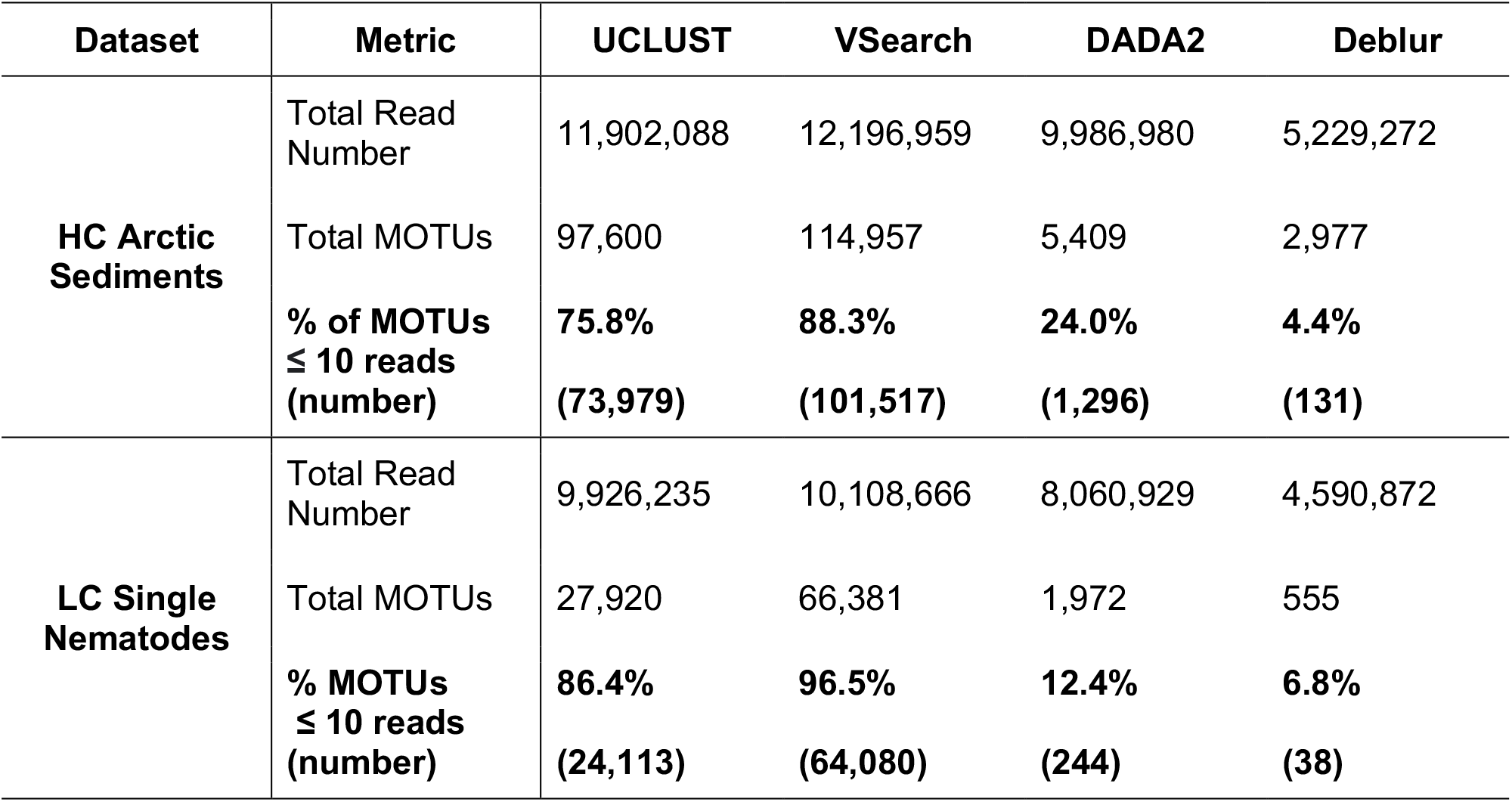
Total number of processed reads, MOTUs, and rare MOTUs (≤10 reads) for each dataset and pipeline. Bold text indicates the percentage and number of rare MOTUs in the HC and LC datasets.

| Dataset | Metric | UCLUST | VSearch | DADA2 | Deblur |
| --- | --- | --- | --- | --- | --- |
| HC Arctic Sediments | Total Read Number | 11,902,088 | 12,196,959 | 9,986,980 | 5,229,272 |
|  | Total MOTUs | 97,600 | 114,957 | 5,409 | 2,977 |
| | % of MOTUs $\leq 10$ reads | **75.8%** | **88.3%** | **24.0%** | **4.4%** |
|  | (number) | **(73,979)** | **(101,517)** | **(1,296)** | **(131)** |
| LC Single Nematodes | Total Read Number | 9,926,235 | 10,108,666 | 8,060,929 | 4,590,872 |
|  | Total MOTUs | 27,920 | 66,381 | 1,972 | 555 |
| | % MOTUs $\leq 10$ reads | **86.4%** | **96.5%** | **12.4%** | **6.8%** |
|  | (number) | **(24,113)** | **(64,080)** | **(244)** | **(38)** |

We also detected a significant positive correlation between the number of MOTUs and retained reads in both the OTU and ASV algorithms, except for Deblur in the LC dataset (R^2^ = 0.01, p-value = 0.31). Although all four pipelines showed this relationship, correlations were more moderate for methods estimating ASVs (Figure 2C, 2F). This correlation between MOTUs and retained reads may be impacted by microbial community complexity and sequencing depth. Furthermore, when looking at subsets of each dataset (i.e., Arctic subregions and nematode families for the HC and LC datasets, respectively; Figure S2, S3), these patterns of recovered reads and MOTUs were highly consistent across bioinformatics pipelines, particularly for the HC dataset. For example, no significant differences (p > 0.05) were detected among Arctic subregions for both MOTUs and retained reads (Figure S2). For the LC dataset, we also observed an overall agreement across algorithms for the number of reads, except for Deblur which differed from the others (e.g., family Oxystomindae; Figure S3D). However, for MOTUs we observed changes (i.e., from non-significant to significant) within and between algorithm classes in the LC dataset (Figure S3).

### MOTU rank abundance and taxonomy profiles

Given the significant differences in total number of reads and MOTUs recovered across algorithm classes (Tables S1, S2), we next sought to investigate the source of this discrepancy and assess the potential implications for biological interpretations of environmental metabarcoding datasets. Rank-abundance curves were generated for MOTUs recovered across all four pipelines in the HC and LC datasets (Figure 4A, 4B). In both datasets, the OTU algorithms (UCLUST and Vsearch) exhibited a typical L-shaped rank abundance curve with a steep gradient, indicating that top ranked OTUs had much higher abundances than the “long tail” of rare OTUs. OTU datasets were almost entirely dominated by rare MOTUs, representing anywhere from 75.8% to 96.5% of the entire dataset (Table 3). In contrast, the ASV algorithms showed S-shaped (DADA2) and C-shaped (Deblur) rank-abundance curves with much more gentle slopes. Overall, these patterns seem not to be impacted by dataset complexity.

**Figure 4.**
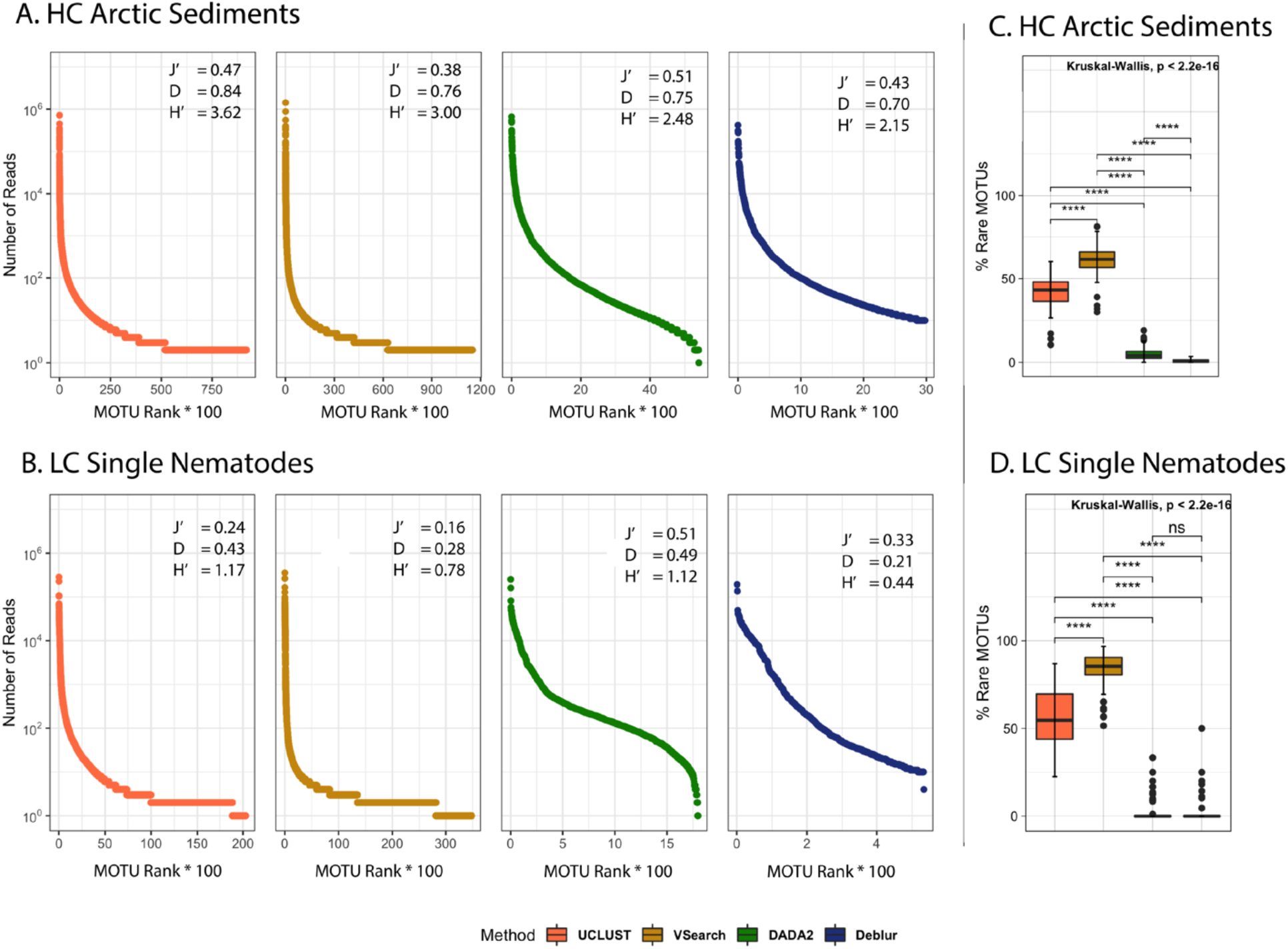
Ranked MOTU distribution showing “Head-Tail” patterns across pipelines. Rank-abundance curves for the HC Arctic sediment (**A**) and LC single nematode (**B**) datasets. In the x-axis, the number of ranked MOTUs is multiplied by a factor of 100. Alpha-diversity indices including Shannon (H’), Simpson (D), and Pielou’s evenness (J’) are also given for each method/dataset. For rare MOTUs (i.e., containing ≤ 10 reads), Kruskal-Wallis analysis was used to test for significant differences among bioinformatics pipelines for HC (**C**) and LC (**D**) datasets.

Both DADA2 and Deblur strongly reduced the collection of rare MOTUs (e.g., the “long tail”) commonly found in metabarcoding datasets (Figure 4, Table 3). The Deblur pipeline returned the lowest proportion of rare MOTUs across both datasets (i.e., only 4.4% and 6.8% of the HC and LC datasets, respectively), while DADA2 exhibited slightly higher proportions of rare MOTUs (24% and 12.4% of OTUs in the HC and LC datasets, respectively). This difference between ASV algorithms is also apparent in the MOTU rank-abundance curves (Deblur having a shorter tail of rare MOTUs; Figure 4). We also noted that dataset complexity appears to impact the percentage of rare MOTUs recovered in ASV pipelines: the HC dataset contained a higher proportion of rare MOTUs in DADA2 (as expected given the presumed higher sample biodiversity), while Deblur returned a higher proportion of rare MOTUs in the LC dataset, suggesting that dataset complexity may be a confounding factor for the Deblur algorithm performance. Overall, there appeared to be a higher evenness amongst MOTUs produced from ASV algorithms (DADA2 and Deblur), which is reflected by the slope of the curves (i.e., gentle slopes indicating less dominance of fewer top ranked ASVs) and the fewer low-ranked ASVs (Figure 4). Moreover, deeper exploration of the LC dataset (i.e., four major nematode families) confirmed that the behavior between methods within the same MOTU class is more similar than when comparing across classes and also that OTU clustering algorithms tend to produce longer low-abundance tails (Figure S5).

Although the shape of rank-abundance curves varied across methods in the HC and LC datasets, the abundance of the dominant MOTU (i.e., the first rank MOTU) was consistent across all pipelines, and about an order of magnitude higher than the second most abundant MOTU (Figure 4). In the LC dataset where we could evaluate the MOTU profile of individual nematodes, the nucleotide sequence of the most abundant (Head) MOTU remained identical across all four algorithms (Table 4). Moreover, we observed that Head MOTUs in ASV pipelines showed a 10-20% reduction in relative abundance, presumably because ASVs are a less inclusive “cloud” of sequence reads. When examining Head MOTUs in the HC dataset, however, congruence was only detected between algorithms of the same MOTU class, which also led to some different taxonomic assignments (Table 4). The primary difference across pipelines remained the curve shape and length of the “long tail” of rare MOTUs, with algorithm class having the strongest influence (OTU vs. ASV pipelines). Similar patterns were also apparent in subgroups of the LC dataset, where OTU algorithms appeared to produce a longer and more prominent tail of rare MOTUs (e.g., Chromadoridae, Figure S5). Interesting, for the highest-ranked MOTUs (i.e., the top 10 most abundant), the different algorithms behaved fairly similar, except for Deblur (family Oxystominidae; Figure S5). Despite these observed changes in MOTU rank-abundance curves, the relative abundance of major taxonomic groups remained fairly stable across all four bioinformatics pipelines in both the LC and HC datasets (Figure 3). Small differences were only observed in the LC dataset, particularly a higher abundance of Comesomatidae in the Deblur method and the absence of some lower abundance taxa, which varied according to pipeline (e.g., Filobasidiaceae in UCLUST outputs; Xyalidae in Vsearch outputs; Linhomoeidae in DADA2 outputs; Microlaimidae in Deblur outputs; Figure 3).

**Table 4.** The top 10 most abundant MOTUs for each bioinformatics pipeline in the HC and LC datasets. UCLUST taxonomy and relative abundance (RA %) is used as a reference for comparison with the other three methods.

| Method | MOTU | RA (%) | HC Dataset – Taxonomy | Vsearch (RA%) | Dada2 (RA%) | Deblur (RA%) |
| --- | --- | --- | --- | --- | --- | --- |
| UCLUST | MOTU1 | 6.0 | Annelida; Polychaeta; Scolecida; Spionida | OTU1 (11.69) | ASV2 (5.37) | ASV2 (5.91) |
|  | MOTU2 | 6.0 | Annelida; Polychaeta; Palpata; Phyllodocida | OTU2 (7.20) | ASV1 (6.64) | ASV1 (7.96) |
|  | MOTU3 | 3.8 | Ochrophyta; Diatomea; Bacillariophytina; Mediophyceae; Chaetoceros | OTU3 (4.5) | ASV4 (4.83) | ASV3 (5.77) |
|  | MOTU4 | 3.0 | Annelida; Polychaeta; Scolecida; Spionida | - | **ASV3 (5.0)** | **ASV4 (5.26)** |
|  | MOTU5 | 2.9 | Nematoda; Chromadorea; Araeolaimida; Comesomatidae; *Sabatieria*; *Sabatieria* sp. | OTU4 (3.25) | ASV6 (2.95) | ASV6 (3.15) |
|  | MOTU6 | 2.7 | Dinoflagellata; Dinophyceae; Peridiniphyctidae; Gonyaulacales; *Alexandrium* | OTU6 (2.78) | ASV5 (3.17) | ASV5 (3.31) |
|  | MOTU7 | 2.5 | Annelida; Polychaeta; Scolecida; Spionida | - | - | - |
|  | MOTU8 | 2.3 | Annelida; Polychaeta; Scolecida; Spionida | OTU7 (2.67) | ASV7 (2.66) | ASV7 (2.90) |
|  | MOTU9 | 1.9 | Annelida; Polychaeta; Scolecida; Spionida | OTU9 (2.05) | ASV8 (2.16) | ASV8 (2.79) |
|  | MOTU10 | 1.8 | Annelida; Polychaeta; Scolecida; Spionida | OTU5 (3.04) | ASV9 (2.15) | ASV9 (2.33) |
| Method | MOTU | RA (%) | Taxonomy – LC Dataset | Vsearch (RA%) | Dada2 (RA%) | Deblur (RA%) |
| UCLUST | MOTU1 | 7.13 | Nematoda; Chromadorea; Chromadorida; Chromadoridae | OTU1 (8.83) | ASV1 (8.01) | ASV1 (12.12) |
|  | MOTU2 | 5.69 | Nematoda; Chromadorea; Araeolaimida; Comesomatidae; *Sabatieria*; *Sabatieria* sp. | OTU2 (6.55) | ASV2 (5.09) | ASV2 (8.46) |
|  | MOTU3 | 2.71 | Nematoda; Chromadorea; Desmodorida | OTU4 (3.15) | ASV3 (2.6) | ASV3 (3.09) |
|  | MOTU4 | 2.59 | Nematoda; Chromadorea; Araeolaimida; Comesomatidae; *Sabatieria*; *Sabatieria* sp. | OTU3 (4.08) | ASV4 (1.84) | ASV4 (2.6) |
|  | MOTU5 | 1.76 | Nematoda; Chromadorea; Araeolaimida; Comesomatidae | OTU9 (1.78) | ASV5 (1.61) | ASV5 (2.45) |
|  | MOTU6 | 1.67 | Nematoda; Enoplea; Enoplida; Oxystominidae | OTU6 (2.16) | ASV9 (1.52) | - |
|  | MOTU7 | 1.62 | Nematoda; Chromadorea; Araeolaimida; Comesomatidae | OTU8 (1.84) | ASV6 (1.56) | ASV6 (2.41) |
|  | MOTU8 | 1.6 | Nematoda; Chromadorea; Desmoscolecida; Desmoscolecidae; *Desmoscolex* | - | ASV10 (1.34) | ASV8 (2.12) |
|  | MOTU9 | 1.59 | Nematoda; Enoplea; Enoplida; Thoracostomopsidae | OTU5 (2.41) | - | ASV10 (1.77) |
|  | MOTU10 | 1.48 | Nematoda; Chromadorea; Araeolaimida; Comesomatidae; *Sabatieria*; *Sabatieria* sp. | OTU7 (1.86) | - | - |

MOTUs that agreed in the taxonomic assignment across pipelines but differed in one nucleotide are highlighted in bold.
(-) It indicates no UCLUST correspondent MOTU was detected for that specific pipeline.

### Alpha-diversity metrics across pipelines

Alpha-diversity metrics were calculated for both HC and LC datasets, using MOTU tables resulting from all four bioinformatics pipelines. To assess fine-scale differences, each metabarcoding study was analyzed according to biologically relevant sample groupings including six Arctic geographic subregions (HC dataset), and four well-sampled nematode families (LC dataset). As observed in the full datasets (Figure 2), the number of OTUs recovered for each HC and LC data subset remained about fifteen-fold higher than the number of ASVs (Figures S2, S3) across all subgroups, indicating that OTU pipelines consistently inflate MOTU diversity at all levels of a metabarcoding dataset. With respect to Shannon and Simpson diversity (Figure S4), there were typically no significant differences among subgroups (i.e., Arctic subregion or nematode family), except for the DADA2 pipeline in the LC dataset. Thus, the same subgroups were consistently recovered as having the lowest and highest median values of alpha-diversity regardless of the MOTU pipeline (e.g., for the HC dataset Shannon Diversity metric, Beaufort Shelf and Camden Bay exhibited the lowest median value, while Bank Islands exhibited the highest median value Figure S4).

Conversely, bioinformatics pipelines did influence the absolute alpha-diversity of each specific subgroup (Figures 5, 6). For example, ASV pipelines resulted in significant reductions of Shannon diversity for the HC Arctic sediments (Figure 5), as opposed to Simpson diversity which was overall more stable across pipelines (although we did observe significant differences across methods for Arctic regions with > 30 samples, Amundsen Gulf, Chukchi Sea, and Mackenzie River Plume; Figure 6). For the LC dataset, Shannon and Simpson indices were more consistent, although bioinformatics pipelines appeared to have disproportionately higher effects on the calculated diversity of each nematode family. Almost all pipeline comparisons resulted in significant differences in alpha-diversity for each LC dataset subgroup (Figures 5, 6). Surprisingly, the difference between UCLUST and DADA2 was rarely significant, with these two algorithms returning similar levels of Shannon and Simpson diversity despite representing different MOTU algorithm classes. Both dataset complexity and the number of samples representing a specific subgroup appeared to be responsible for the significant differences observed in these diversity indices.

**Figure 5.**
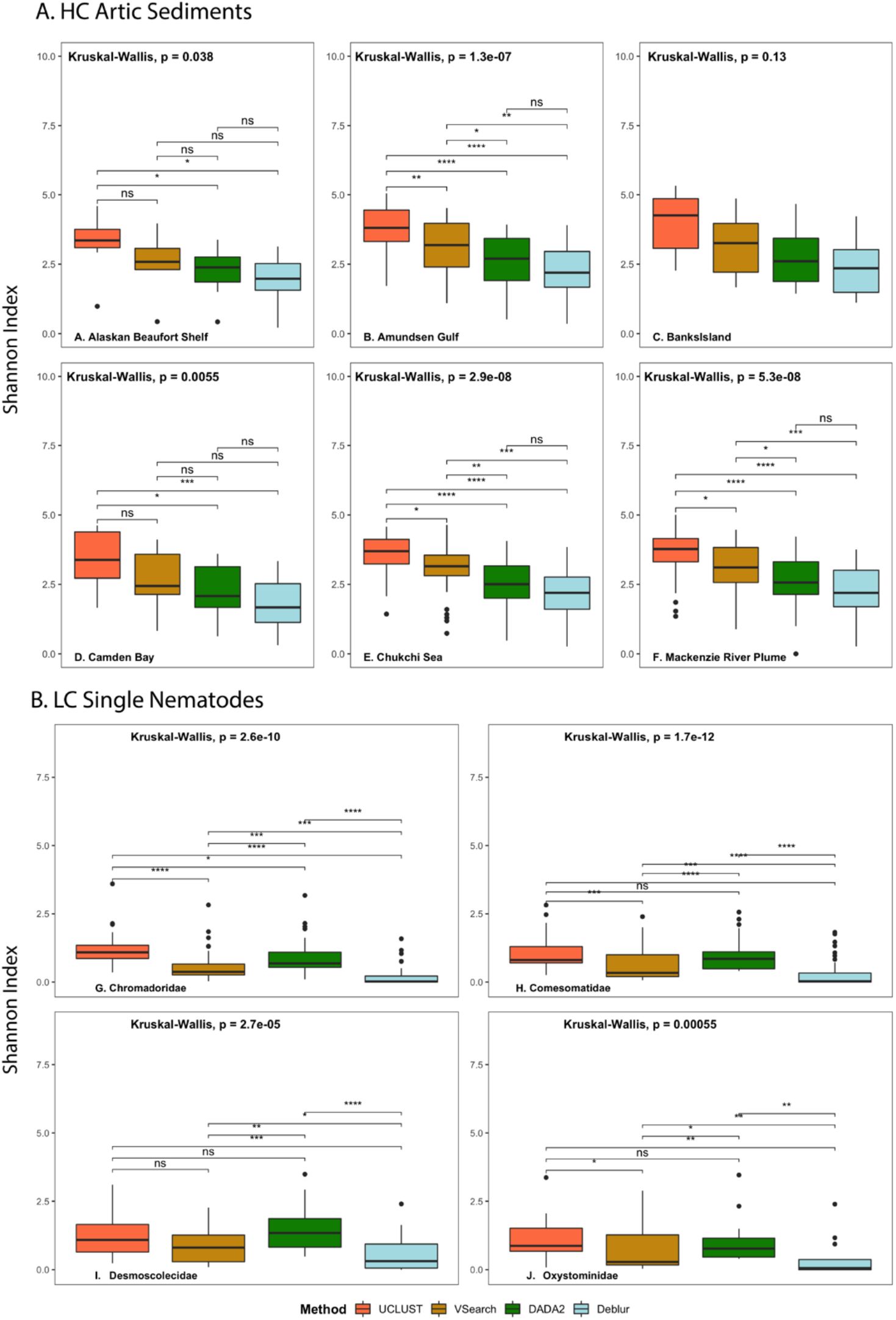
Shannon diversity index for (A) HC and (B) LC datasets. Kruskal-Wallis analysis was used to test for significant differences among bioinformatics pipelines for each subregion (HC dataset) or nematode family (LC dataset), separately.

**Figure 6.**
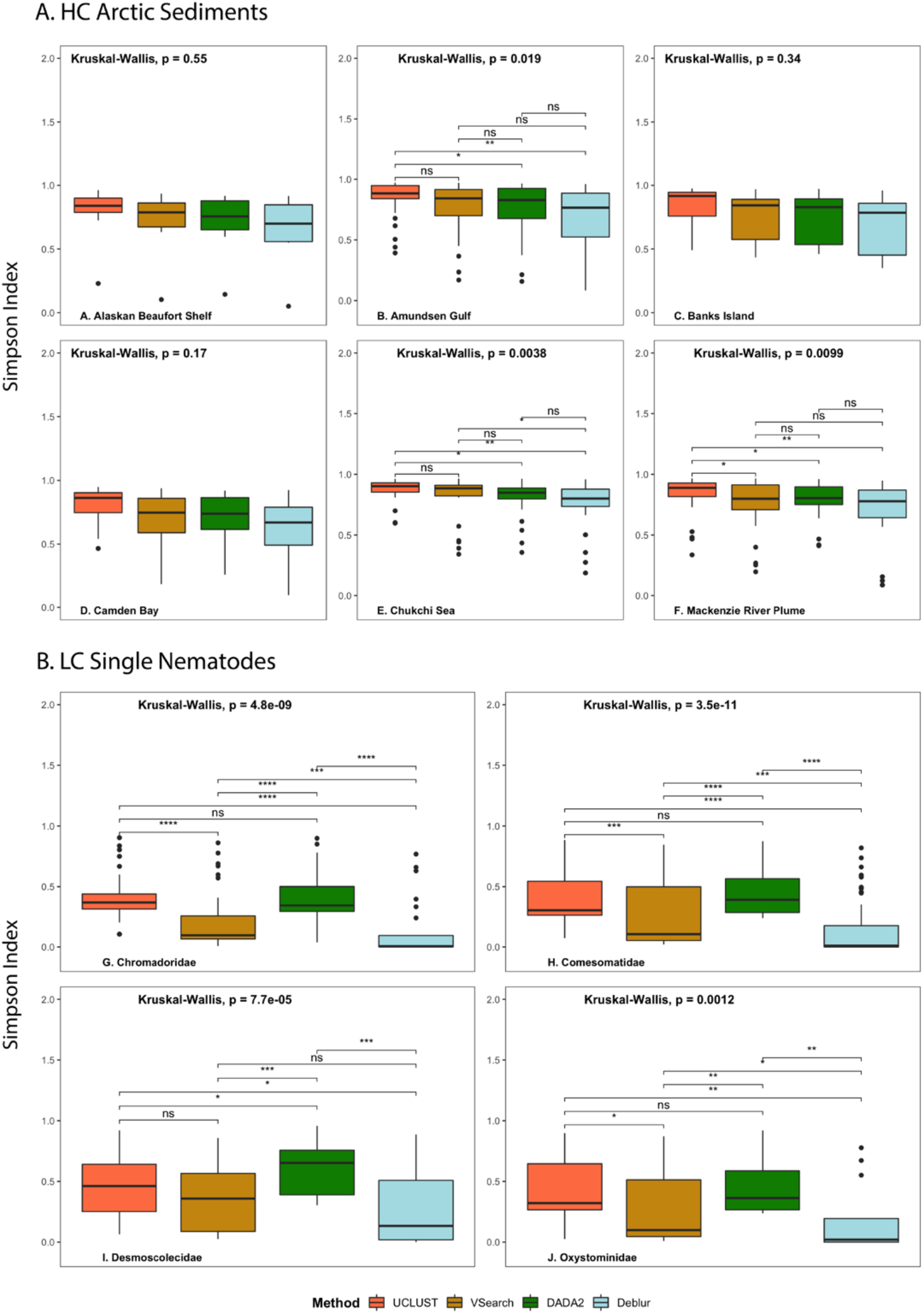
Simpson diversity index for (A) HC and (B) LC datasets. Kruskal-Wallis analysis was used to test for significant differences among bioinformatics pipelines for each subregion (HC dataset) or nematode family (LC dataset), separately.

### Beta-diversity metrics across pipelines

Beta-diversity metrics were also calculated for both HC and LC datasets using results from all four pipelines and based on the Bray-Curtis, unweighted Unifrac, and weighted Unifrac indices (rarefied at 1000 sequences per sample). The Procrustes analysis always revealed significant concordance (p < 0.01) between pipelines regardless of the dataset, sampling depth, or distance index (Table 2, Figure 7). Our results suggest that beta-diversity metrics, as well as dataset complexity, can both impact the level of concordance observed in beta-diversity analyses across bioinformatics pipelines. For both the HC and LC datasets, Bray-Curtis similarity exhibited high concordance across all pairwise comparisons of bioinformatics pipelines, especially between DADA2 and Deblur. Apparently, neither algorithm class (OTU vs. ASV generation) nor dataset complexity (HC vs. LC dataset) impacted the beta-diversity patterns recovered with the Bray-Curtis metric as observed by the relatively similar statistical values and the length of vectors connecting data points (Table 2, Figure 7). In contrast, dataset complexity had a clear influence on the level of concordance observed for both weighted and unweighted Unifrac metrics. In the LC dataset, most pairwise Unifrac comparisons exhibited low concordance (e.g., R <0.8 and M^12^ >0.2; Table 2), except for UCLUST-Vsearch (unweighted Unifrac), VSearch-DADA2, and DADA2-Deblur (both weighted Unifrac) which exhibited high concordance. In contrast, all pairwise comparisons across both weighted/unweighted Unifrac metrics were recovered as highly concordant in the HC dataset.

**Figure 7.**
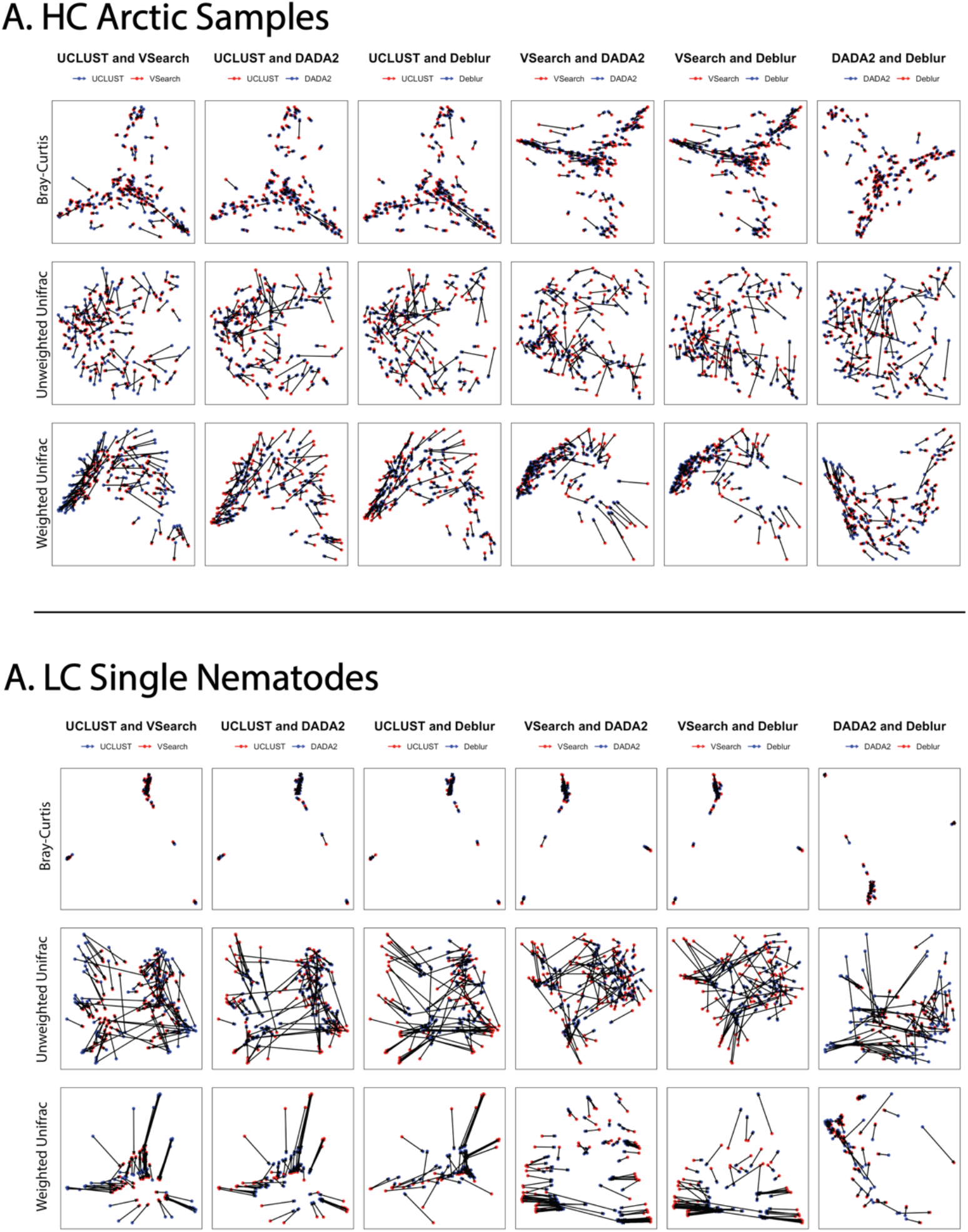
Procrustes Analysis for beta-diversity. A Procrustes ordination plot based on PCoA coordinates was made for each beta diversity index (e.g., Bray-Curtis, unweighted Unifrac, and weighted Unifrac). Pairwise comparisons between bioinformatics pipelines were showed the highest congruence when using the Bray-Curtis. The colors underneath each method refers to their color on the plot. Longer lines connecting samples indicate higher discordance in beta-diversity patterns, whereas short lines (or no visible lines) denote the highest concordance.

## Discussion

This study illustrates the differential influence of four bioinformatics pipelines on eukaryotic 18S rRNA metabarcoding studies. We focused on UCLUST, VSearch, DADA2, and Deblur since these workflows represent the most commonly applied approaches in the 18S rRNA metabarcoding literature. We did not attempt an exhaustive assessment of all software tools that exist for the delimitation of MOTUs (Boyer et al., 2016; Edgar, 2016; Eren et al., 2015; Mahé et al., 2014; Schloss et al., 2009). However, most MOTU algorithms process metabarcoding reads in ways that are conceptually similar to our four chosen workflows, and thus our results should be broadly generalizable across studies. Algorithm class (OTU vs. ASV delimitation) appeared to be the strongest driver of the observed differences in both the HC and LC datasets, having a clear impact on both the number of quality-processed reads and MOTUs (Figure 2), the shape of MOTU rank-abundance curves (Figures 4A, 4B), and the proportion of “rare biosphere” taxa represented by low abundance MOTUs (Figures 4C, 4D, Table 3). However, dataset complexity also impacted bioinformatics outputs in subtle and unexpected ways. For example, LC datasets appeared to introduce an element of randomness and stochasticity into the downstream ecological analysis (e.g., beta-diversity metrics; Table 2, Figure 7), and the specific influence of algorithm parameters became more extreme for metabarcoding samples with simpler community structure (e.g., patterns of recovered reads and MOTUs for the LC dataset; Table 3). In fact, Siegwald et al. (2017) pointed out that the analysis of low complexity datasets is often a difficult task, perhaps because it contains a higher number of low abundance taxa (i.e., rare MOTUs).

In our LC dataset (where per-sample biodiversity consisted of one nematode and a small number of host-associated taxa and gut contents; Pereira et al., 2020), the VSearch algorithm returned 2.4x as many OTUs compared to UCLUST despite these two pipelines being in the same MOTU class (Table 3). Furthermore, we observed that the “long tail” of rare MOTUs was even more pronounced in the VSearch pipeline, especially for the LC dataset (i.e., 96.5% of all OTUs). These trends appear to be related to the underlying functions of the two OTU algorithms. While UCLUST utilizes a heuristic method to find read alignments with the best score (via implementation of USEARCH; Edgar, 2010), the VSearch algorithm instead generates the full alignment vectors during MOTU generation (via global pairwise alignments using the Needleman-Wunsch algorithm; Rognes et al., 2016). Therefore, it is possible that VSearch produces tighter clusters (i.e., with smaller sequence divergence) leading to a higher number of OTUs, including more rare MOTUs. Previous benchmark studies have demonstrated that UCLUST is likely to produce looser clusters, especially at lower similarity thresholds (Ghodsi et al., 2011). Despite the higher proportion of rare MOTUs in VSearch outputs, previous studies have indicated that VSearch exhibits equal or higher accuracy and returns more stable OTU clusters (that are unlikely to disappear when the data is subsetted or increased) compared to the UCLUST algorithm when performing *de novo* clustering (Jackson et al., 2016; Westcott & Schloss, 2015). The “long tail” of rare MOTUs appears to be an inherent feature of OTU picking pipelines where sequences are clustered via pairwise sequence identity. However, our results suggest that the relative proportion of rare MOTUs (and the dominance of this “long tail” in any given metabarcoding dataset) will vary according to algorithm choice as well as the underlying dataset complexity, thus supporting previous findings (Siegwald et al., 2017, 2019).

In ASV approaches, the Deblur algorithm also appeared to be especially sensitive to dataset complexity. In our LC dataset, the contribution of rare MOTUs was higher than expected in the Deblur pipeline (6.8% of ASVs, which was ∼1.5x more than in the HC dataset; Table 3). Similarly, read error correction was markedly more conservative in our LC dataset, where Deblur showed a significant reduction in the number of ASVs recovered compared to DADA2 (∼15% reduction; Figure 2D). In contrast, the difference between the two ASV pipelines was not significant in the HC dataset, with Deblur returning a higher number of MOTUs than DADA2; Figure 2A). However, Deblur always returned a lower *<u>total</u>* number of reads and MOTUs for the overall HC and LC datasets compared to DADA2 (Tables S1, S2). For mock communities, Prodan et al. (2020) reported that Deblur conserved only about 50% of the initial read number input, whereas other pipelines were above 70%. Accordingly, the authors related Deblur’s low conversion rate to the count-subtraction nature of the algorithm before the ASV estimation. This discrepancy between the total vs. mean number of recovered reads/MOTUs appears to stem from the fundamentally distinct ways in which DADA2 and Deblur perform quality checks and error correction on raw Illumina data (Amir et al., 2017; Callahan et al., 2016). DADA2 merges raw Illumina reads after its ASV-generation algorithm (i.e., after trimming, truncating, and denoising), while Deblur merges reads as a first step before subsequent quality checks are performed. Most importantly, Deblur performs all error correction and ASV generation steps on a sample-by-sample basis. The per-sample focus was a conscious choice by the Deblur developers, who designed this ASV algorithm to maximize computational speed and maintain the ability to be highly parallelizable when required for large datasets. However, this software design can erroneously remove sequences considered as “rare MOTUs” by only viewing slices of a large HTS dataset. DADA2 instead generates an error-correction model by considering all samples and sequences at once (e.g., with the underlying assumption that each Illumina run has a unique profile of sequencing artefacts that can be eliminated by comparing abundant vs. rare reads). As a result, DADA2 is much more computationally intensive and requires more computational time. Our results suggest that the inherent features of the Deblur algorithm produce less consistent outputs that can be heavily influenced by the biological community complexity built into a metabarcoding study, and potentially impacting the ability of statistical tests to detect significant differences between groups (e.g., subregions and nematode families in our study). Existing ASV algorithms show wide divergence in algorithm design and function, and therefore must be chosen with care.

We observed a surprising level of stability for some biological patterns recovered in downstream ecological analyses. For example, the relative abundance of major taxonomic groups is generally preserved across bioinformatics pipelines regardless of dataset complexity (Figure 3). Although MOTU clustering algorithm did lead to some significant differences in alpha-diversity comparisons (Figures 5, 6), these did not appear to impact biological interpretations within a specific dataset and method, that is, the most diverse subgroup (Arctic subregion or nematode family) was rarely affected in Simpson and Shannon diversity metrics (Figure S4). A previous study by Jackson et al. (2016) confirmed that the absolute values of alpha-diversity indices cannot be compared across bioinformatics methods, and such differences in diversity estimates can be effectively eliminated by collapsing MOTUs according to their taxonomy assignments. However, we note that MOTU taxonomy assignments will also be impacted by different bioinformatics methods (e.g., using BLAST searches vs. Bayesian classifier tools) as well as the completeness of reference databases (Holovachov et al., 2017; Macheriotou et al., 2019), and both of these factors typically vary depending on the chosen metabarcoding workflow and target taxon. When assessing the beta-diversity among subgroups based on the Bray-Curtis coefficient, we also found high concordance across all methods in the HC and LC datasets (Figure 7), presumably because the Bray-Curtis coefficient as a resemblance measure has valid underlying conditions (e.g., independence of joint absence) and tends to capture the important assemblage relationships (Clarke et al., 2006). Taken together, our results indicate that many overarching biological patterns within 18S rRNA metabarcoding datasets remain consistent across distinct bioinformatics pipelines, regardless of dataset complexity. Therefore, beta-diversity patterns appear to be largely unaffected by the overall number of MOTUs (Table 3) or the number of rare MOTUs as depicted in the Head-Tail curves (Figure 4).

Downstream analyses that took rare MOTUs into account were notably less consistent across the four pipelines, especially for the LC dataset. Procrustes analysis of beta-diversity patterns revealed low concordance across bioinformatics pipelines for the LC dataset when the Unifrac distance was applied (Table 2). Since Unifrac relies on the topology of a phylogenetic tree to calculate the evolutionary distance of MOTUs between samples (Lozupone & Knight, 2005), a low number of MOTUs with skewed relative abundance profiles may heavily impact beta-diversity analyses across bioinformatics pipelines (e.g., a few dominant MOTUs containing the vast majority of reads). We believe that LC datasets may be especially sensitive to sample rarefaction employed before Unifrac calculations, which potentially increases the stochasticity of the MOTUs being subsampled and makes beta-diversity comparisons more prone to random effects. From a biological perspective, ASV pipelines offer significant advantages for downstream ecological analyses where rarefaction is employed as they tend to be more stable regardless of the rarefaction depth (Prodan et al., 2020). Furthermore, they strongly reduce the likelihood that sequencing artefacts and erroneous reads will be incorporated into diversity metrics that emphasize changes in the “rare biosphere” of low-read MOTUs. For alpha-diversity specifically, other methods may also be considered when comparing sample groups (e.g., Distanced which use mean-pairwise distance (MPD) using a Bayesian approach; Hackmann, 2020).

For metabarcoding studies, sequencing technologies and bioinformatics pipelines will inevitably continue to evolve. Our prime consideration was to assess the stability of biological inferences across historical (and future) shifts in computational workflows for MOTU generation. For published studies relying on cluster-based OTU methods, the recent shift from QIIME1 (where UCLUST is the default algorithm; Caporaso et al., 2010) to QIIME2 (where VSearch is the default algorithm; Bolyen et al., 2019) is not likely to have a significant impact on the downstream biological conclusions within this class of algorithm. OTU pipelines report similar (albeit highly inflated) levels of biodiversity, and results from UCLUST and VSearch were generally consistent regardless of study design and dataset complexity (HC vs LC datasets; Figures 2, 4, 5, 6). Major patterns of taxon abundance (Figure 3) and sample grouping (ordinations based on Bray-Curtis coefficient; Figure 7) were also unaffected by algorithm class. With the evolution of new algorithms in recent years, the metabarcoding and microbial ecology communities have rapidly shifted towards ASVs as a more stable and objective type of MOTU where reference sequences can be directly compared across studies (Callahan et al., 2017). OTUs are somewhat arbitrary “clouds’’ of sequence reads, and OTU membership can be heavily influenced by the specific parameters of the underlying algorithm (He et al., 2015). Furthermore, OTUs are dataset-specific and not directly comparable across studies (Callahan et al., 2017). Our results confirm the significant biological advantages of ASV generation algorithms: the improved error correction eliminates the artefactual “long tail” of rare sequences while maintaining species-specific barcodes (“Head” MOTUs with high relative abundance; Porazinska, Giblin-Davis, Esquivel, et al., 2010). More importantly, ASVs do not completely eliminate rare MOTUs, and the remaining low-abundance reads can provide important biological insights regarding ecological interactions and population-level variation (e.g., patterns of intragenomic rRNA variation (Pereira et al., 2020; Qing et al., 2020), gut contents and host-associated microbiome taxa (Schuelke et al., 2018)). We specifically recommend the DADA2 pipeline for eukaryotic metabarcoding studies, as the resulting ASV dataset appears to represent the best approximation of real biological patterns (and the number of distinct species present), especially for LC communities where the sequencing effort has likely reached saturation (Macheriotou et al., 2019; Pereira et al., 2020). The Deblur algorithm should be used with caution, however, since Deblur’s optimization for fast computational speed has resulted in a tradeoff whereby biologically valid MOTUs (true positives) are overzealously eliminated from metabarcoding datasets. Further analysis of already published 18S rRNA metabarcoding studies using the DADA2 pipeline may reveal compelling ecological and evolutionary insights for diverse eukaryotic taxa on a global scale given the increased biological accuracy of this method (e.g., shorter tail of rare MOTUs, less likely to be impacted by data complexity).

The present study focused on a metabarcoding locus optimized for microbial metazoa, targeting the V1-V2 regions of the 18S rRNA gene (Creer et al., 2010). It remains to be seen whether the observed bioinformatics patterns will extend to other rRNA loci and protein-coding genes. For soil nematodes specifically, Kenmotsu et al. (2020) suggested that the amplification of regions V7-V9 (located at 3’ end of the 18S rRNA gene) followed by the analysis using DADA2 may produce the most realistic results. Future research efforts should evaluate computational pipelines using other common metabarcoding genes (COI, 12S, rbcl, etc.) and 18S rRNA hypervariable regions (V4, V9, etc.) thus incorporating broader assessments of both nuclear and mitochondrial loci (e.g, PEMA pipeline; Zafeiropoulos et al., 2020). How dataset complexity influences bioinformatics outputs is likely to be distinct for metabarcoding loci such as COI, where Illumina datasets will inherently contain a high level of both population-level (e.g., gene haplotypes) and species-level (“Head” MOTUs) genetic variation. Although DADA2 performed well with both HC and LC datasets, and therefore should be the chosen method for 18S rRNA metabarcoding studies, our study also supports the long-held assumption that major biological and ecological patterns will tend to emerge from a well-designed metabarcoding study (Xiong & Zhan, 2018; Zinger et al., 2019), regardless of the underlying bioinformatics pipeline used for data analysis. Summarizing MOTU datasets by taxonomy assignments (e.g., collapsing MOTUs at the genus or species level) appears to be a robust approach for avoiding any discrepancies that may arise across different computational pipelines (Jackson et al., 2016), and this approach may help to alleviate some of the observed difficulties in analyzing rare MOTUs and LC datasets. However, such a taxonomy-dependent approach requires a comprehensive database of reference DNA barcodes (Ruppert et al., 2019). Unfortunately, current reference databases for common eukaryotic metabarcoding loci are comparatively sparse and patchy (versus 16S rRNA gene databases for bacteria/archaea where most known genera are well represented; Bik et al., 2012; Macheriotou et al., 2019; Pereira et al., 2020). Improvement of eukaryotic reference databases, combined with a movement towards phylogeny-based biodiversity analyses, will help to further improve the ecological and evolutionary metrics that can be applied to metabarcoding studies.

## Data Availability Statement

Raw Illumina reads from Arctic marine sediments (HC dataset) are deposited in the NCBI Sequence Read Archive (BioProject PRJNA723923 and SRA accession SUB9498354). For the Low Complexity (LC) single-nematode dataset, raw Illumina reads are also available in the NCBI Sequence Read Archive (BioProject PRJNA422296 and SRA accession SRP128131). Additionally, full-length 18S rRNA gene sequences were generated via Sanger sequencing for most nematode specimens used in the LC dataset and deposited on GenBank (Accession Numbers: MN250033–MN250142). Primer constructs for both HC and LC dataset have been deposited in FigShare (https://doi.org/10.6084/m9.figshare.5701090). QIIME mapping files, final MOTU tables, and all scripts used for processing and analyzing the data are available on GitHub (https://github.com/BikLab/OTU-ASV-euk-benchmarking).

## Author Contributions

ADS, TJP, and HMB conceived the ideas and designed study methodology; SMH collected marine sediment samples and provided intellectual input on hypothesis testing and data analysis workflows; ADS collated and analyzed the data; ADS, TJP, and HMB led the writing of the manuscript. All authors contributed critically to the drafts and gave final approval for publication.

## Acknowledgments

The authors would like to thank Taruna Schuelke and Alexis Walker for their help and assistance in generating the original metabarcoding datasets used as the basis for this study. Funding for this study was provided by the North Pacific Research Board (NPRB project 1303), The Gulf of Mexico Research Initiative, and institutional startup funding from the University of Georgia. Data are publicly available through the Gulf of Mexico Research Initiative Information & Data Cooperative (GRIIDC) at https://data.gulfresearchinitiative.org (doi: R5.x272.000:0004).

---

**Figure S1.**
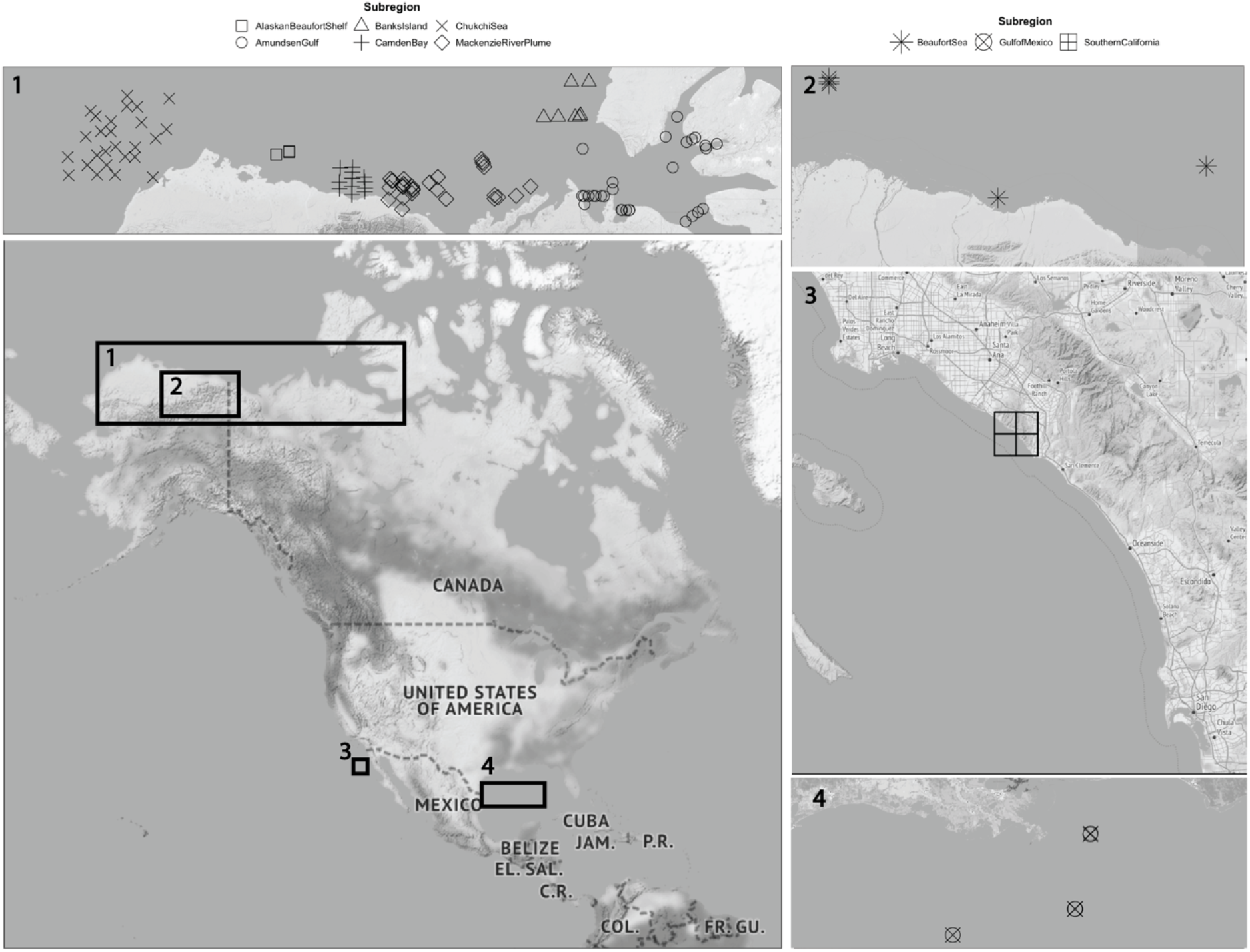
Geographical distribution of samples used in this study. Samples from the “high complexity” (HC) dataset were collected from the Alaskan Arctic region and included six subregions. Single nematodes from the “low complexity” (LC) dataset were isolated from sediment cores collected from the Alaskan Arctic, Gulf of Mexico, and Southern California (an intertidal holdfast site) regions.

**Figure S2.**
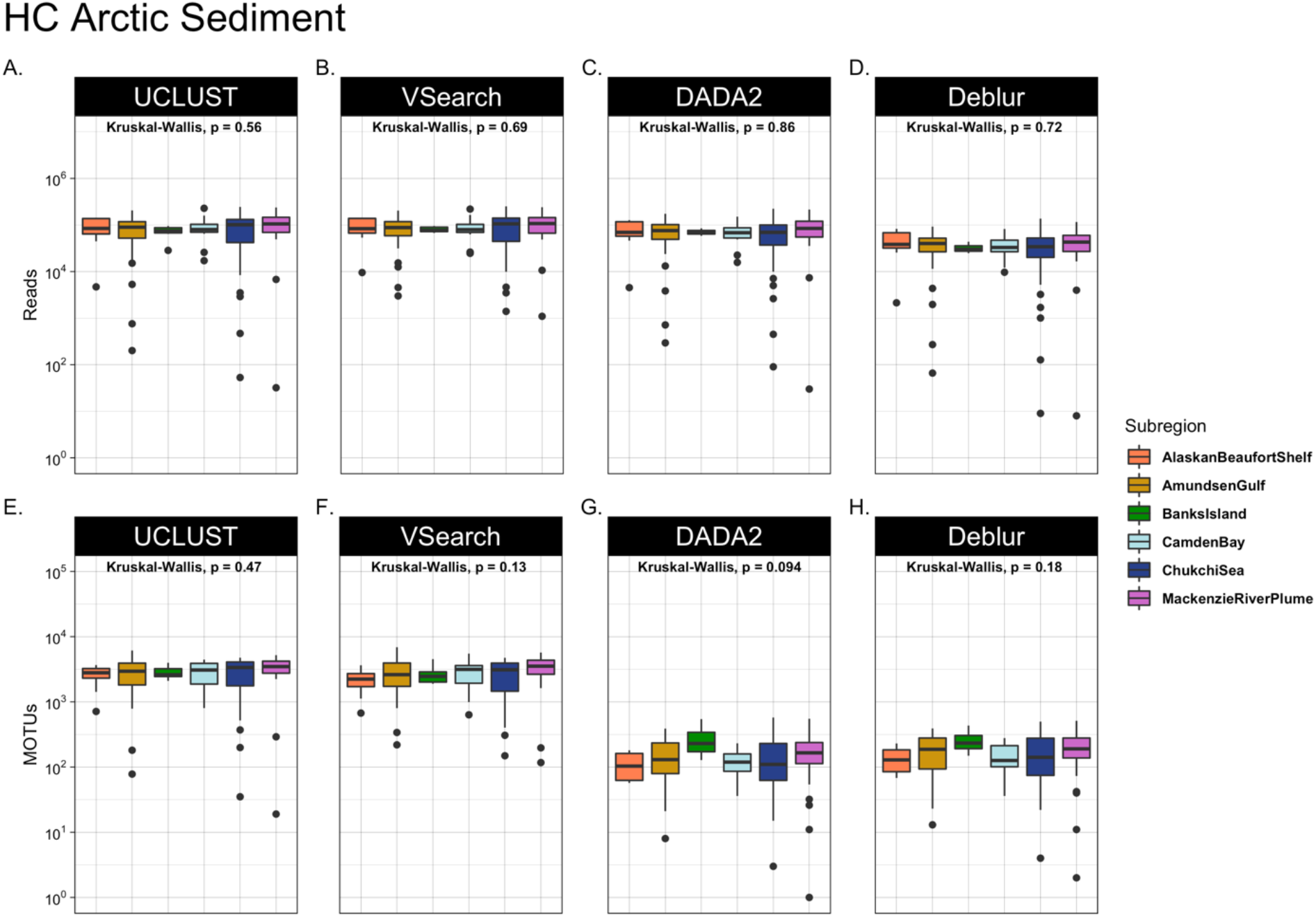
Number of reads (A-D) and MOTUs (E-H) for each Arctic subregion in the HC dataset. Kruskal-Wallis analysis was used to test for significant differences among subregions for each bioinformatic pipeline, separately.

**Figure S3.**
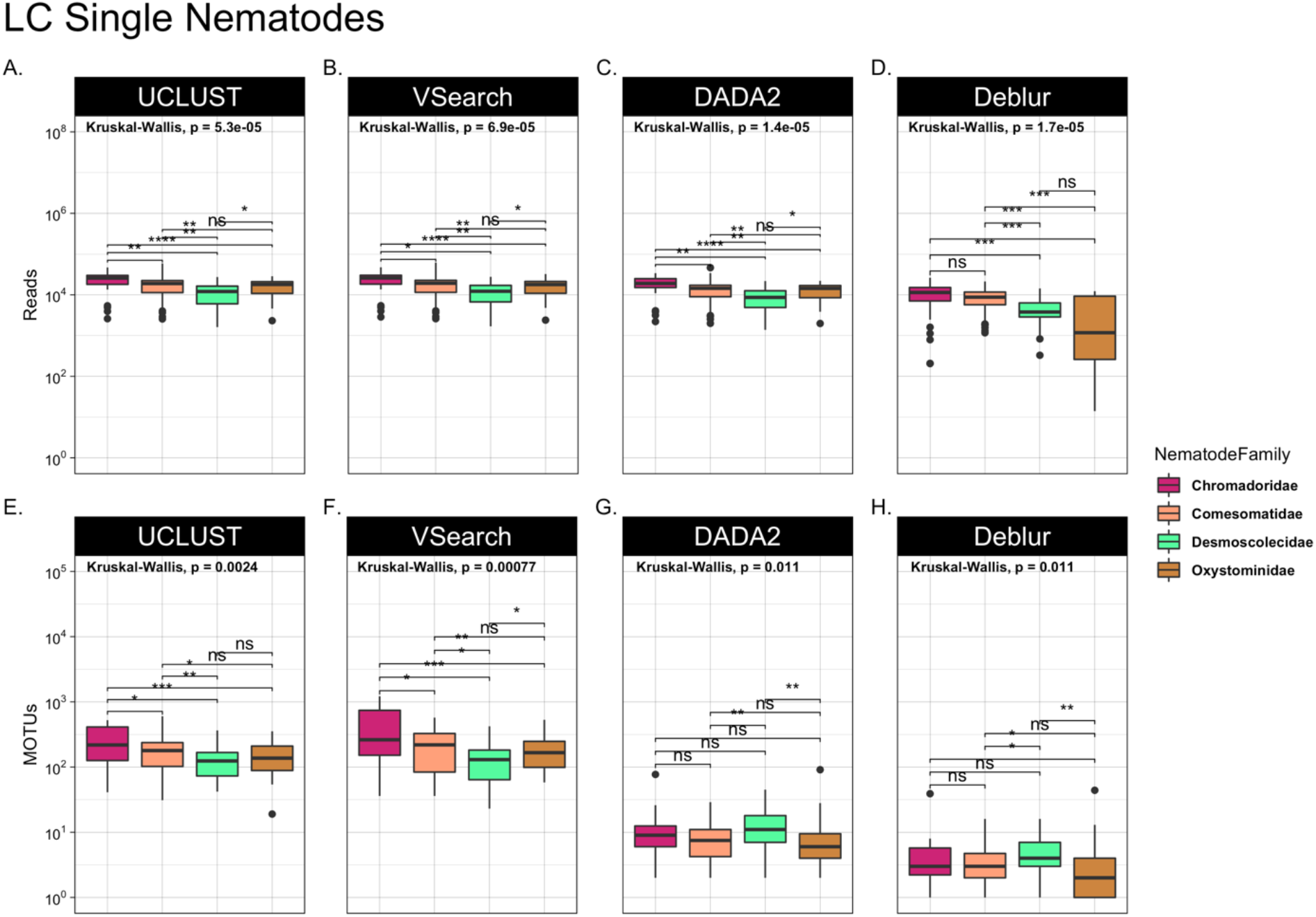
Number of reads (A-D) and MOTUs (E-H) for four major nematode families in the LC dataset. Kruskal-Wallis analysis was used to test for significant differences among nematode families for each bioinformatic pipeline, separately.

**Figure S4.**
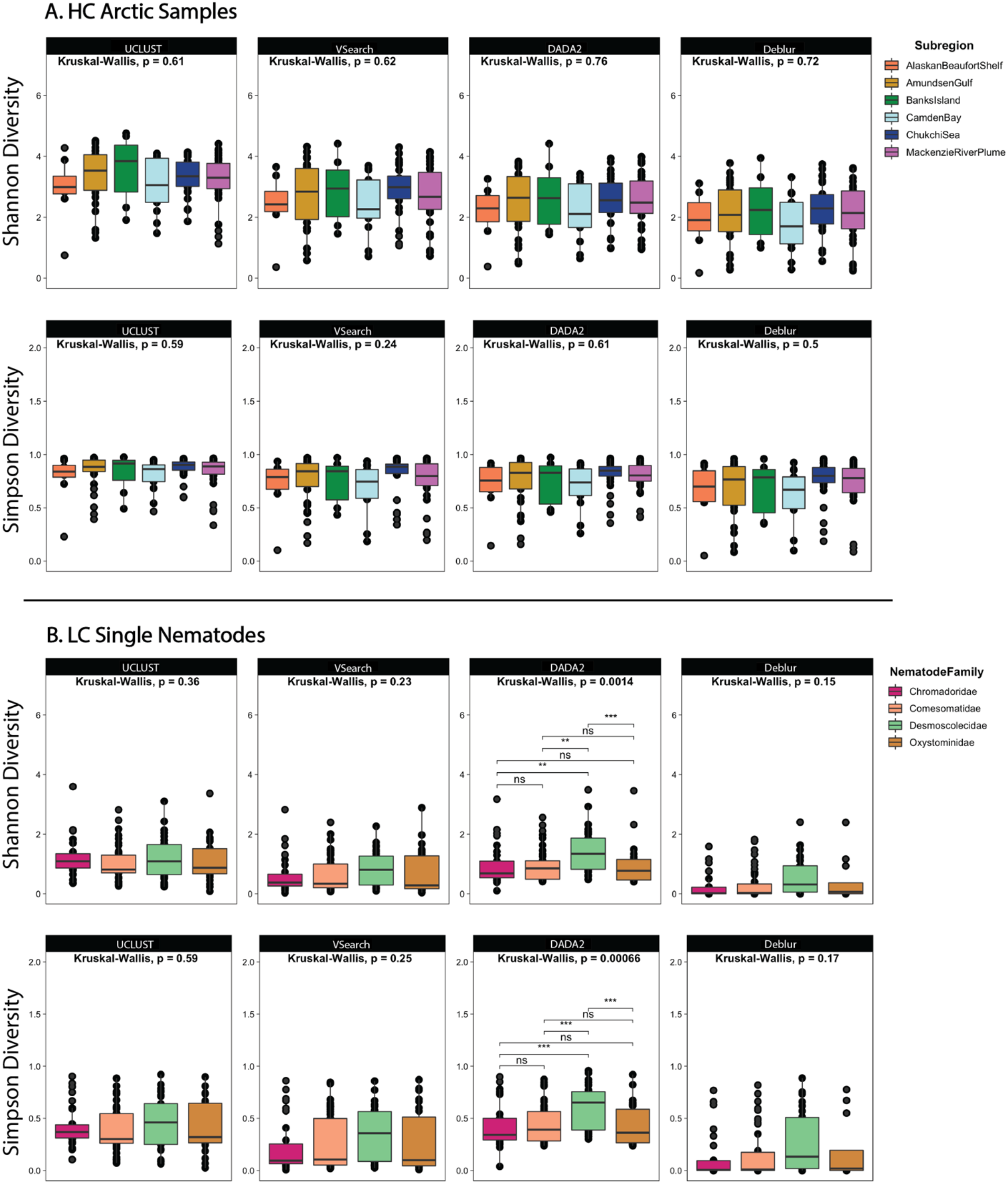
Shannon and Simpson diversity indices for (A) HC and (B) LC datasets. Kruskal-Wallis analysis was used to test for significant differences among subregions (HC dataset) and nematode families (LC dataset) for each bioinformatics pipeline, separately.

**Figure S5.**
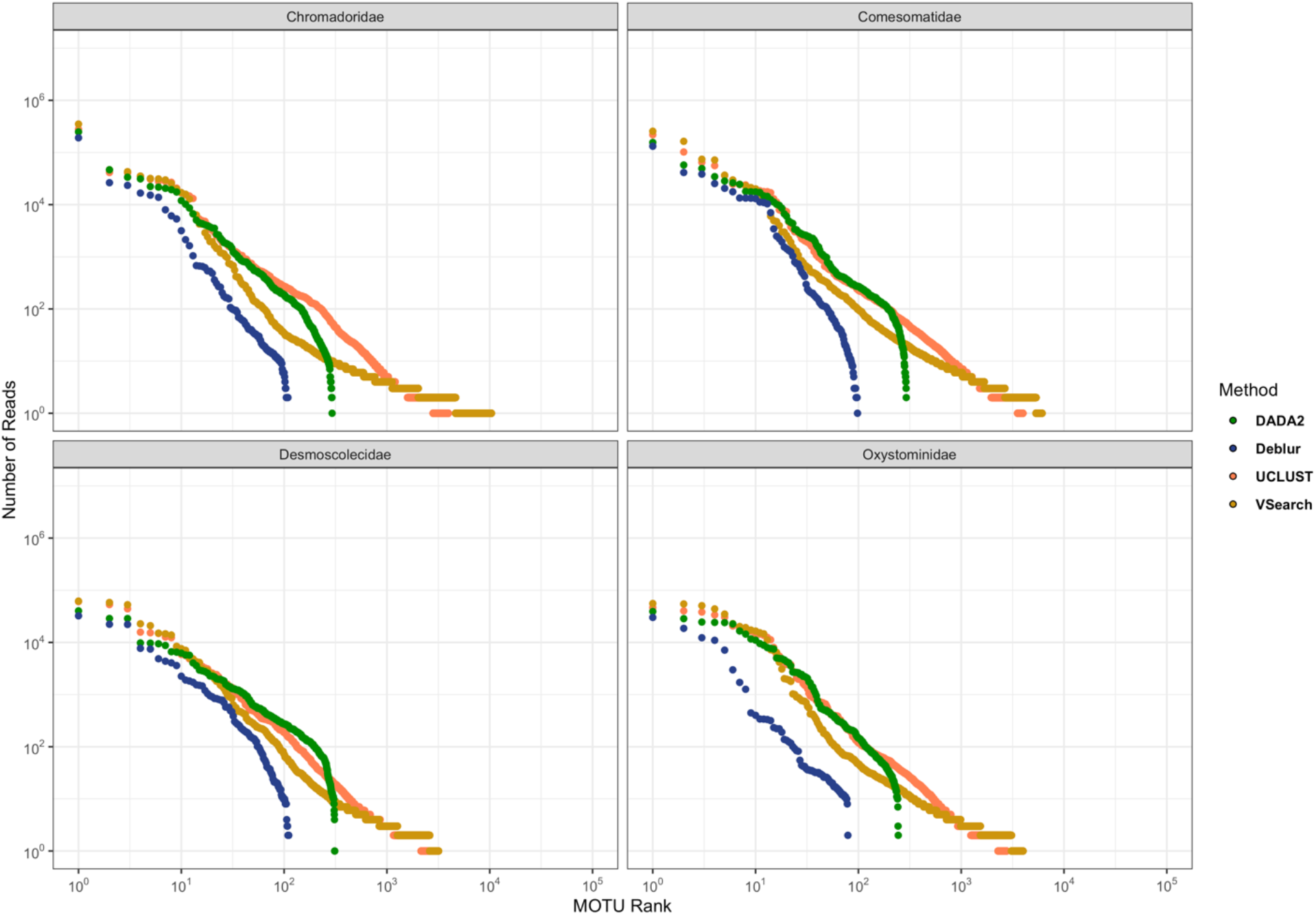
Ranked MOTU distribution showing “Head-Tail” patterns across the different pipelines for four major nematode families represented in the LC dataset.

