## Supplementary material for "Dataset complexity impacts both MOTU delimitation and biodiversity estimates in eukaryotic 18S rRNA metabarcoding studies": Table S2

| **Samples** | **UCLUST** | | **VSearch** | | **DADA2** | | **Deblur** | |
| --- | --- | --- | --- | --- | --- | --- | --- | --- |
|  | **Reads** | **OTUs** | **Reads** | **OTUs** | **Reads** | **ASVs** | **Reads** | **ASVs** |
| **Nem17** | 20,985 | 180 | 21,121 | 197 | 15,918 | 25 | 9,975 | 13 |
| **Nem18** | 22,048 | 391 | 22,446 | 672 | 19,007 | 12 | 11,792 | 3 |
| **Nem19** | 20,492 | 66 | 20,580 | 101 | 16,908 | 6 | 10,969 | 1 |
| **Nem20** | 25,029 | 154 | 25,187 | 189 | 16,966 | 14 | 11,573 | 4 |
| **Nem21** | 31,381 | 482 | 31,738 | 963 | 26,654 | 10 | 17,175 | 3 |
| **Nem22** | **14,262** | 354 | 14,471 | 382 | 10,989 | 91 | 3,678 | 44 |
| **Nem23** | 5,390 | 218 | 5,422 | 162 | 4,080 | 77 | 792 | 39 |
| **Nem24** | 20,929 | 307 | 21,256 | 306 | 17,577 | 26 | 10,598 | 14 |
| **Nem25** | 18,086 | 182 | 18,299 | 255 | 15,248 | 2 | 9,864 | 1 |
| **Nem26** | 29,278 | 185 | 30,168 | 286 | 23,268 | 5 | 13,770 | 1 |
| **Nem27** | 23,038 | 74 | 23,124 | 94 | 19,037 | 3 | 12,294 | 1 |
| **Nem28** | 18,844 | 160 | 18,966 | 182 | 15,446 | 22 | 9,441 | 11 |
| **Nem29** | 11,766 | 143 | 11,793 | 149 | 8,346 | 26 | 4,419 | 13 |
| **Nem30** | 28,915 | 466 | 29,274 | 887 | 24,167 | 13 | 15,144 | 6 |
| **Nem31** | 15,450 | 286 | 16,368 | 394 | 9,620 | 21 | 4,114 | 9 |
| **Nem32** | 21,411 | 168 | 21,965 | 177 | 17,273 | 5 | 11,144 | 4 |
| **Nem33** | 13,765 | 167 | 13,902 | 221 | 11,586 | 4 | 7,509 | 3 |
| **Nem34** | 22,517 | 175 | 22,677 | 125 | 18,076 | 11 | 11,362 | 5 |
| **Nem35** | 17,024 | 145 | 17,177 | 166 | 13,245 | 8 | 6,506 | 4 |
| **Nem36** | 37,161 | 326 | 37,993 | 798 | 25,465 | 10 | 9,177 | 5 |
| **Nem37** | 21,780 | 270 | 22,053 | 393 | 18,186 | 20 | 11,702 | 13 |
| **Nem38** | 23,126 | 217 | 23,664 | 327 | 15,985 | 8 | 9,404 | 3 |
| **Nem39** | 19,007 | 359 | 19,234 | 579 | 16,158 | 8 | 10,767 | 3 |
| **Nem40** | 4,320 | 73 | 4,580 | 71 | 3,109 | 8 | 2,122 | 7 |
| **Nem41** | 19,422 | 300 | 19,682 | 530 | 15,381 | 8 | 3,963 | 4 |
| **Nem42** | 27,938 | 105 | 27,992 | 115 | 22,402 | 5 | 14,641 | 3 |
| **Nem43** | 16,227 | 364 | 17,147 | 278 | 12,297 | 24 | 5,921 | 16 |
| **Nem44** | 16,357 | 305 | 16,997 | 296 | 7,830 | 45 | 4,489 | 14 |
| **Nem45** | 15,073 | 84 | 15,293 | 70 | 12,602 | 15 | 7,848 | 11 |
| **Nem46** | 9,915 | 124 | 10,106 | 130 | 7,611 | 14 | 3,414 | 3 |
| **Nem47** | 20,823 | 372 | 21,097 | 679 | 17,819 | 9 | 11,068 | 3 |
| **Nem48** | 17,008 | 365 | 17,198 | 561 | 14,485 | 12 | 9,187 | 5 |
| **Nem49** | 29,979 | 90 | 30,126 | 152 | 23,813 | 19 | 206 | 8 |
| **Nem50** | 6,031 | 103 | 6,070 | 67 | 4,887 | 11 | 2,864 | 5 |
| **Nem51** | 20,285 | 233 | 20,709 | 390 | 15,844 | 6 | 7,535 | 3 |
| **Nem52** | 17,334 | 85 | 17,364 | 89 | 14,197 | 9 | 260 | 4 |
| **Nem53** | 17,397 | 132 | 17,579 | 151 | 13,718 | 12 | 7,310 | 6 |
| **Nem54** | 17,400 | 130 | 17,602 | 127 | 12,986 | 29 | 6,040 | 11 |
| **Nem55** | 25,028 | 124 | 25,549 | 166 | 20,453 | 13 | 411 | 4 |
| **Nem56** | 7,494 | 124 | 8,079 | 175 | 6,020 | 25 | 3,369 | 16 |
| **Nem57** | 7,429 | 92 | 7,710 | 79 | 5,787 | 25 | 843 | 12 |
| **Nem58** | 19,702 | 226 | 20,124 | 330 | 14,342 | 17 | 4,841 | 10 |
| **Nem59** | 28,640 | 214 | 28,908 | 310 | 24,156 | 3 | 15,682 | 5 |
| **Nem60** | 2,895 | 66 | 2,945 | 60 | 2,249 | 17 | 441 | 15 |
| **Nem61** | 5,445 | 83 | 5,554 | 103 | 4,281 | 18 | 1,622 | 8 |
| **Nem63** | 19,556 | 86 | 20,159 | 88 | 16,618 | 5 | 9,865 | 1 |
| **Nem64** | 3,642 | 102 | 3,672 | 68 | 2,971 | 25 | 1,171 | 16 |
| **Nem65** | 27,683 | 270 | 28,102 | 343 | 20,052 | 11 | 14,440 | 3 |
| **Nem66** | 29,727 | 235 | 30,584 | 329 | 23,249 | 15 | 13,869 | 3 |
| **Nem67** | 21,681 | 248 | 21,999 | 282 | 15,734 | 2 | 11,318 | 1 |
| **Nem68** | 14,581 | 335 | 14,812 | 524 | 12,381 | 8 | 7,280 | 2 |
| **Nem69** | 22,198 | 229 | 22,534 | 290 | 15,911 | 8 | 11,755 | 3 |
| **Nem70** | 23,894 | 265 | 24,286 | 344 | 16,958 | 2 | 12,164 | 5 |
| **Nem71** | 19,117 | 163 | 19,295 | 226 | 16,008 | 5 | 10,109 | 3 |
| **Nem72** | 18,574 | 231 | 18,849 | 228 | 13,405 | 2 | 9,216 | 2 |
| **Nem73** | 16,008 | 147 | 16,160 | 198 | 13,698 | 4 | 8,726 | 2 |
| **Nem74** | 24,764 | 316 | 25,221 | 339 | 17,855 | 7 | 12,373 | 5 |
| **Nem75** | 11,011 | 147 | 11,166 | 212 | 8,987 | 11 | 5,598 | 7 |
| **Nem76** | 18,126 | 104 | 18,199 | 128 | 15,129 | 7 | 9,512 | 2 |
| **Nem77** | 11,787 | 188 | 12,025 | 213 | 9,667 | 6 | 4,982 | 2 |
| **Nem78** | 13,298 | 197 | 13,488 | 199 | 9,475 | 3 | 6,801 | 1 |
| **Nem79** | 22,432 | 238 | 22,747 | 289 | 16,078 | 3 | 11,766 | 2 |
| **Nem80** | 16,003 | 140 | 16,302 | 173 | 13,600 | 12 | 8,024 | 7 |
| **Nem81** | 18,008 | 117 | 17,921 | 157 | 14,943 | 10 | 9,276 | 4 |
| **Nem84** | 2,058 | 43 | 2,102 | 36 | 1,658 | 6 | 939 | 3 |
| **Nem85** | 5,867 | 123 | 5,932 | 69 | 4,598 | 10 | 2,494 | 5 |
| **Nem86** | 15,560 | 163 | 15,993 | 170 | 12,359 | 18 | 4,823 | 4 |
| **Nem87** | 23,717 | 173 | 24,077 | 203 | 19,075 | 18 | 12,126 | 9 |
| **Nem88** | 12,055 | 108 | 12,228 | 116 | 9,874 | 7 | 6,331 | 3 |
| **Nem90** | 34,846 | 322 | 35,741 | 803 | 23,999 | 14 | 10,273 | 2 |
| **Nem91** | 46,871 | 176 | 47,378 | 262 | 34,096 | 3 | 26,341 | 1 |
| **Nem92** | 36,851 | 239 | 37,591 | 322 | 31,802 | 4 | 20,285 | 3 |
| **Nem93** | 31,069 | 232 | 31,729 | 309 | 26,926 | 5 | 16,866 | 2 |
| **Nem94** | 31,466 | 157 | 31,750 | 239 | 22,640 | 13 | 15,461 | 8 |
| **Nem95** | 57 | 5 | 96 | 16 | 42 | 2 | 9 | 2 |
| **Nem96** | 1,753 | 36 | 1,775 | 19 | 1,280 | 5 | 568 | 3 |
| **Nem97** | 4,915 | 45 | 4,950 | 45 | 3,830 | 7 | 2,459 | 4 |
| **Nem98** | 791 | 33 | 870 | 45 | 583 | 10 | 332 | 10 |
| **Nem99** | 50 | 9 | 60 | 15 | 35 | 2 | 27 | 2 |
| **Nem100** | 703 | 22 | 727 | 32 | 565 | 7 | 405 | 8 |
| **Nem101** | 3,313 | 95 | 3,346 | 36 | 2,773 | 10 | 1,527 | 7 |
| **Nem103** | 8,575 | 73 | 8,639 | 69 | 5,911 | 13 | 3,409 | 7 |
| **Nem105** | 219 | 30 | 288 | 25 | 229 | 6 | 88 | 5 |
| **Nem106** | 17,868 | 163 | 17,983 | 87 | 14,398 | 10 | 7,570 | 5 |
| **Nem108** | 4,892 | 52 | 4,962 | 64 | 3,796 | 10 | 1,901 | 4 |
| **Nem109** | 20,812 | 200 | 21,435 | 270 | 15,740 | 21 | 6,387 | 3 |
| **Nem110** | 4,212 | 105 | 3,468 | 86 | 3,124 | 11 | 1,929 | 7 |
| **Nem111** | 1,679 | 51 | 1,756 | 56 | 1,365 | 8 | 563 | 5 |
| **Nem112** | 866 | 8 | 911 | 20 | 696 | 5 | 373 | 3 |
| **Nem113** | 23,629 | 47 | 23,665 | 74 | 18,557 | 6 | 10,345 | 3 |
| **Nem114** | 17,649 | 303 | 18,415 | 771 | 14,904 | 14 | 8,233 | 4 |
| **Nem115** | 8,609 | 31 | 8,637 | 29 | 6,679 | 5 | 4,566 | 3 |
| **Nem116** | 51,678 | 220 | 52,091 | 360 | 42,261 | 21 | 27,061 | 19 |
| **Nem117** | 16,646 | 103 | 16,848 | 102 | 13,081 | 11 | 7,357 | 8 |
| **Nem118** | 274 | 21 | 286 | 20 | 215 | 2 | 154 | 2 |
| **Nem119** | 5,459 | 74 | 5,523 | 90 | 4,383 | 16 | 1,589 | 10 |
| **Nem125** | 872 | 37 | 962 | 43 | 718 | 4 | 218 | 1 |
| **Nem126** | 1,346 | 74 | 1,368 | 59 | 1,106 | 10 | 445 | 5 |
| **Nem127** | 1,485 | 32 | 1,744 | 37 | 1,310 | 7 | 647 | 4 |
| **Nem128** | 4,487 | 53 | 4,506 | 80 | 3,650 | 12 | 1,569 | 8 |
| **Nem129** | 886,794 | 3,062 | 913,273 | 8744 | 683,251 | 62 | 333,403 | 7 |
| **Nem130** | 8,772 | 72 | 8,825 | 62 | 7,149 | 10 | 4,207 | 7 |
| **Nem131** | 5,087 | 231 | 5,279 | 243 | 4,137 | 24 | 1,659 | 10 |
| **Nem132** | 23,498 | 169 | 23,666 | 116 | 18,586 | 8 | 12,201 | 5 |
| **Nem133** | 22,643 | 136 | 22,738 | 156 | 18,217 | 15 | 9,819 | 4 |
| **Nem134** | 14,686 | 179 | 12,887 | 137 | 10,433 | 17 | 5,408 | 10 |
| **Nem135** | 12,585 | 131 | 10,542 | 105 | 9,389 | 7 | 5,014 | 4 |
| **Nem136** | 1,612 | 44 | 1,688 | 39 | 1,382 | 7 | 829 | 6 |
| **Nem137** | 11,898 | 208 | 12,712 | 264 | 10,238 | 23 | 3,167 | 7 |
| **Nem138** | 24,079 | 114 | 24,111 | 122 | 20,076 | 7 | 67 | 3 |
| **Nem139** | 34,163 | 188 | 34,428 | 248 | 27,866 | 12 | 16,249 | 3 |
| **Nem140** | 24,104 | 258 | 24,537 | 311 | 20,296 | 4 | 12,531 | 2 |
| **Nem141** | 4,528,706 | 5,185 | 4,581,763 | 27218 | 3,788,697 | 50 | 2,474,314 | 5 |
| **Nem142** | 21,596 | 364 | 21,836 | 647 | 18,479 | 6 | 12,259 | 3 |
| **Nem143** | 20,669 | 130 | 20,945 | 230 | 17,366 | 11 | 10,506 | 2 |
| **Nem144** | 12,381 | 173 | 12,558 | 175 | 9,024 | 3 | 6,641 | 2 |
| **Nem145** | 28,082 | 437 | 28,440 | 942 | 24,191 | 8 | 14,178 | 2 |
| **Nem146** | 35,142 | 523 | 35,656 | 1215 | 29,336 | 11 | 14,953 | 1 |
| **Nem147** | 32,385 | 386 | 32,673 | 367 | 24,940 | 14 | 12,779 | 4 |
| **Nem148** | 18,054 | 175 | 18,230 | 193 | 13,623 | 11 | 8,189 | 4 |
| **Nem149** | 29,963 | 439 | 30,315 | 895 | 25,678 | 6 | 16,844 | 1 |
| **Nem150** | 554 | 36 | 571 | 28 | 452 | 4 | 230 | 3 |
| **Nem151** | 21,540 | 109 | 21,643 | 162 | 15,692 | 6 | 11,556 | 3 |
| **Nem152** | 6,439 | 61 | 6,243 | 69 | 5,255 | 8 | 3,152 | 3 |
| **Nem153** | 12,688 | 243 | 12,832 | 179 | 9,950 | 15 | 4,819 | 6 |
| **Nem154** | 28,238 | 433 | 28,572 | 808 | 23,792 | 15 | 15,188 | 8 |
| **Nem155** | 4,219 | 55 | 4,242 | 56 | 3,687 | 2 | 2,365 | 1 |
| **Nem156** | 22,555 | 168 | 22,907 | 188 | 17,960 | 25 | 1,564 | 10 |
| **Nem157** | 28,665 | 259 | 32,418 | 377 | 21,339 | 28 | 0 | 0 |
| **Nem158** | 20,488 | 284 | 20,651 | 251 | 16,230 | 17 | 8,383 | 3 |
| **Nem159** | 24,410 | 99 | 25,033 | 73 | 20,385 | 4 | 13,463 | 3 |
| **Nem160** | 29,616 | 77 | 29,610 | 88 | 25,816 | 2 | 0 | 0 |
| **Nem161** | 4,249 | 119 | 4,376 | 72 | 3,497 | 8 | 1,290 | 4 |
| **Nem162** | 31,929 | 419 | 32,387 | 417 | 25,307 | 26 | 12,470 | 4 |
| **Nem163** | 15,778 | 71 | 15,726 | 70 | 12,658 | 6 | 8,019 | 2 |
| **Nem164** | 15,358 | 103 | 15,497 | 185 | 12,652 | 5 | 7,105 | 1 |
| **Nem165** | 33,698 | 454 | 34,035 | 945 | 28,764 | 6 | 19,111 | 2 |
| **Nem166** | 21,289 | 163 | 21,593 | 156 | 16,679 | 7 | 8,070 | 6 |
| **Nem167** | 16,996 | 70 | 17,057 | 81 | 11,357 | 6 | 7,786 | 4 |
| **Nem168** | 18,335 | 333 | 18,523 | 234 | 14,595 | 16 | 7,646 | 12 |
| **Nem169** | 25,727 | 148 | 25,909 | 151 | 21,211 | 5 | 13,858 | 2 |
| **Nem170** | 23,055 | 332 | 23,281 | 325 | 18,168 | 23 | 8,352 | 4 |
| **Nem171** | 31,803 | 379 | 32,089 | 377 | 25,648 | 11 | 13,614 | 4 |
| **Nem172** | 21,588 | 218 | 21,971 | 298 | 18,185 | 5 | 0 | 0 |
| **Nem173** | 19,842 | 115 | 19,993 | 163 | 17,005 | 2 | 0 | 0 |
| **Nem174** | 4,026 | 73 | 4,054 | 59 | 3,496 | 5 | 1,886 | 2 |
| **Nem175** | 4,733 | 101 | 4,988 | 107 | 3,748 | 16 | 1,319 | 2 |
| **Nem177** | 26,224 | 172 | 26,676 | 272 | 22,074 | 12 | 13,041 | 2 |
| **Nem178** | 33,797 | 191 | 34,289 | 213 | 28,557 | 15 | 15,952 | 5 |
| **Nem179** | 21,999 | 168 | 22,179 | 213 | 16,443 | 12 | 343 | 2 |
| **Nem181** | 25,759 | 187 | 26,058 | 236 | 20,041 | 21 | 86 | 2 |
| **Nem183** | 5 | 5 | 4 | 4 | 0 | 0 | 0 | 0 |
| **Nem184** | 983 | 16 | 1,020 | 31 | 813 | 3 | 479 | 2 |
| **Nem185** | 32,152 | 383 | 33,409 | 964 | 27,493 | 17 | 15,343 | 4 |
| **Nem186** | 15,076 | 121 | 15,514 | 255 | 11,263 | 16 | 4,863 | 8 |
| **Nem187** | 40,655 | 270 | 41,314 | 423 | 34,264 | 13 | 20,695 | 4 |
| **Nem188** | 32,996 | 57 | 33,073 | 116 | 26,477 | 6 | 14,292 | 2 |
| **Nem189** | 19,850 | 118 | 20,179 | 128 | 15,699 | 12 | 7,003 | 4 |
| **Nem190** | 43,194 | 301 | 44,003 | 371 | 34,654 | 11 | 19,731 | 2 |
| **Nem191** | 58,985 | 601 | 61,937 | 573 | 46,394 | 23 | 21,305 | 6 |
| **Nem192** | 5,160 | 226 | 5,334 | 285 | 4,619 | 10 | 2,298 | 6 |
| **Nem193** | 4,956 | 31 | 5,027 | 36 | 3,595 | 6 | 1,932 | 1 |
| **Nem194** | 24,015 | 197 | 24,239 | 190 | 16,914 | 15 | 221 | 1 |
| **Nem195** | 23,168 | 262 | 24,512 | 389 | 19,785 | 13 | 11,014 | 2 |
| **Nem196** | 4,185 | 51 | 4,302 | 37 | 3,226 | 4 | 1,907 | 2 |
| **Nem197** | 8,105 | 9 | 8,135 | 26 | 6,287 | 4 | 4,184 | 1 |
| **Nem200** | 2,596 | 105 | 2,864 | 36 | 2,207 | 5 | 1,606 | 4 |
| **Nem201** | 3,941 | 41 | 3,998 | 38 | 3,150 | 12 | 1,126 | 6 |
| **Nem203** | 1,017 | 25 | 1,052 | 34 | 720 | 2 | 129 | 1 |
| **Nem204** | 14 | 12 | 15 | 13 | 0 | 0 | 0 | 0 |
| **Nem205** | 12,715 | 109 | 12,782 | 111 | 9,394 | 3 | 6,989 | 1 |
| **Nem206** | 17,185 | 115 | 17,400 | 206 | 14,347 | 5 | 8,996 | 1 |
| **Nem207** | 16,749 | 117 | 16,834 | 119 | 12,386 | 4 | 9,607 | 1 |
| **Nem208** | 26,365 | 127 | 26,647 | 254 | 22,228 | 4 | 14,355 | 3 |
| **Nem209** | 14,548 | 208 | 14,907 | 290 | 10,836 | 4 | 5,812 | 2 |
| **Nem210** | 13,644 | **51** | 13,711 | 69 | 10,944 | 4 | 5,100 | 2 |
| **Nem211** | 22,956 | 73 | 23,059 | 84 | 18,679 | 3 | 12,397 | 1 |
| **Nem212** | 21,303 | 125 | 21,441 | 112 | 17,189 | 19 | 208 | 5 |
| **Nem213** | 16,775 | 217 | 17,127 | 288 | 12,263 | 4 | 6,605 | 2 |
| **Nem214** | 24,472 | 84 | 24,564 | 144 | 17,479 | 12 | 7,728 | 2 |
| **Nem215** | 26,766 | 247 | 27,255 | 231 | 22,071 | 6 | 0 | 0 |
| **Nem217** | 20,826 | 259 | 21,309 | 456 | 16,630 | 14 | 7,700 | 3 |
| **Nem219** | 19,926 | 122 | 20,092 | 192 | 16,583 | 9 | 10,289 | 4 |
| **Nem220** | 5,186 | 48 | 5,207 | 48 | 4,196 | 11 | 1,808 | 5 |
| **Nem221** | 19,250 | 190 | 19,659 | 321 | 15,697 | 11 | 9,414 | 6 |
| **Nem222** | 2,556 | 60 | 2,594 | 42 | 1,996 | 5 | 1,315 | 2 |
| **Nem224** | 19,966 | 238 | 20,852 | 383 | 15,870 | 14 | 8,081 | 4 |
| **Nem225** | 2,901 | 43 | 2,915 | 42 | 2,527 | 4 | 1,572 | 3 |
| **Nem227** | 725 | 23 | 738 | 30 | 590 | 1 | 398 | 1 |
| **Nem228** | 33,140 | 484 | 33,570 | 1035 | 28,473 | 9 | 17,346 | 3 |
| **Nem229** | 4,596 | 68 | 4,788 | 100 | 3,850 | 5 | 302 | 1 |
| **Nem230** | 4,253 | 73 | 4,289 | 51 | 3,223 | 5 | 2,291 | 4 |
| **Nem231** | 23,348 | 119 | 23,448 | 150 | 18,859 | 6 | 11,925 | 3 |
| **Nem232** | 17,984 | 262 | 18,361 | 394 | 14,035 | 22 | 6,062 | 6 |
| **Nem233** | 4,751 | 137 | 4,785 | 214 | 4,066 | 7 | 86 | 1 |
| **Nem234** | 3,005 | 48 | 3,017 | 23 | 2,531 | 2 | 1,603 | 1 |
| **Nem235** | 544 | 17 | 550 | 20 | 486 | 2 | 224 | 1 |
| **Nem237** | 9,237 | 137 | 9,376 | 83 | 7,725 | 2 | 0 | 0 |
| **Nem238** | 2,317 | 56 | 2,396 | 61 | 1,982 | 2 | 1,156 | 1 |
| **Nem239** | 8,795 | 93 | 8,941 | 67 | 6,356 | 8 | 4,014 | 6 |
| **Nem240** | 5,669 | 92 | 5,764 | 98 | 4,656 | 2 | 0 | 0 |
| **Nem241** | 15,343 | 164 | 15,670 | 225 | 11,658 | 39 | 3,555 | 5 |
| **Nem242** | 140,993 | 610 | 142,789 | 1142 | 118,597 | 6 | 73,975 | 2 |
| **Nem243** | 7,447 | 54 | 7,579 | 76 | 6,197 | 4 | 40 | 2 |
| **Nem244** | 5,370 | 52 | 6,438 | 61 | 4,480 | 4 | 2,740 | 1 |
| **Nem245** | 7,946 | 95 | 8,101 | 92 | 6,407 | 8 | 2,895 | 1 |
| **Nem246** | 17,003 | 248 | 17,136 | 451 | 14,400 | 4 | 0 | 0 |
| **Nem247** | 27,110 | 289 | 27,937 | 503 | 18,453 | 26 | 7,500 | 7 |
| **Nem248** | 10,834 | 78 | 11,429 | 89 | 8,215 | 10 | 4,930 | 6 |
| **Nem249** | 14,123 | 155 | 14,213 | 344 | 11,175 | 15 | 5,060 | 4 |
| **Nem250** | 16,521 | 172 | 16,963 | 266 | 11,964 | 9 | 6,009 | 6 |
| **Nem254** | 19,364 | 167 | 19,650 | 134 | 14,163 | 18 | 8,931 | 8 |
| **Nem255** | 19,119 | 139 | 18,726 | 213 | 15,020 | 16 | 4,435 | 5 |
| **Nem256** | 5,265 | 75 | 6,703 | 48 | 5,228 | 11 | 3,170 | 4 |
| **Nem257** | 6,676 | 44 | 6,779 | 64 | 4,880 | 7 | 3,013 | 5 |
| **Nem258** | 20,411 | 122 | 20,633 | 149 | 14,349 | 8 | 7,341 | 3 |
| **Nem260** | 1,750 | 11 | 1,753 | 3 | 1,480 | 2 | 0 | 0 |
| **Nem261** | 2,694 | 135 | 2,742 | 74 | 2,183 | 4 | 1,399 | 2 |
| **Nem262** | 4 | 4 | 5 | 5 | 0 | 0 | 0 | 0 |
| **Nem263** | 16,087 | 208 | 16,966 | 244 | 13,772 | 7 | 8,618 | 2 |
| **Nem264** | 14,676 | 185 | 14,906 | 194 | 10,709 | 5 | 7,550 | 2 |
| **Nem265** | 17,296 | 201 | 17,569 | 227 | 12,702 | 3 | 8,798 | 1 |
| **Nem267** | 18,092 | 210 | 18,392 | 226 | 13,152 | 4 | 9,291 | 1 |
| **Nem268** | 12,147 | 109 | 12,259 | 91 | 9,758 | 5 | 5,591 | 1 |
| **Nem269** | 14,466 | 241 | 14,885 | 233 | 8,980 | 10 | 4,227 | 1 |
| **Nem270** | 19,174 | 164 | 19,425 | 214 | 15,658 | 6 | 8,670 | 1 |
| **Nem271** | 11,422 | 19 | 11,463 | 58 | 9,146 | 6 | 14 | 1 |
| **Nem272** | 15,416 | 221 | 15,948 | 388 | 12,157 | 16 | 311 | 2 |
| **Nem273** | 10,163 | 182 | 10,376 | 167 | 8,641 | 16 | 2,931 | 3 |
| **Nem274** | 2,386 | 58 | 2,465 | 32 | 1,843 | 3 | 1,316 | 2 |
| **Nem275** | 17,289 | 130 | 17,389 | 118 | 13,978 | 8 | 7,946 | 2 |
| **Nem276** | 13,193 | 205 | 13,470 | 279 | 10,274 | 15 | 1,721 | 2 |
| **Nem277** | 22,981 | 269 | 23,679 | 422 | 18,086 | 28 | 329 | 2 |
| **Nem278** | 23,507 | 127 | 23,615 | 163 | 18,298 | 8 | 11,186 | 2 |
| **Nem279** | 12,860 | 133 | 13,482 | 147 | 10,568 | 15 | 5,296 | 5 |
| **Nem280** | 19,491 | 202 | 20,228 | 241 | 16,309 | 10 | 26 | 1 |
| **Nem281** | 336,438 | 1,499 | 344,261 | 3340 | 255,702 | 39 | 78,131 | 2 |
| **Nem282** | 16,665 | 140 | 17,084 | 143 | 13,900 | 9 | 7,321 | 2 |
| **Nem283** | 7,457 | 42 | 7,481 | 43 | 5,559 | 6 | 3,793 | 3 |
| **Nem284** | 347 | 9 | 347 | 5 | 298 | 2 | 187 | 1 |
| **Nem285** | 20,836 | 267 | 21,178 | 331 | 14,930 | 8 | 9,469 | 2 |
| **Nem286** | 11,861 | 166 | 12,052 | 237 | 9,288 | 6 | 338 | 1 |
| **Nem287** | 13,475 | 209 | 13,629 | 134 | 10,927 | 7 | 6,200 | 3 |
| **Nem288** | 10,292 | 124 | 10,430 | 140 | 7,904 | 9 | 1,177 | 4 |
| **Nem289** | 26,281 | 204 | 26,566 | 237 | 21,273 | 10 | 13,370 | 3 |
| **Nem290** | 30,433 | 136 | 30,726 | 197 | 24,462 | 10 | 13,286 | 2 |
| **Nem291** | 27,372 | 185 | 27,673 | 226 | 21,806 | 7 | 14,335 | 4 |
| **Nem292** | 17,897 | 162 | 18,071 | 201 | 13,664 | 9 | 7,393 | 3 |
| **Nem293** | 0 | 0 | 216 | 13 | 0 | 0 | 0 | 0 |
| **Nem294** | 1,779 | 36 | 1,818 | 41 | 1,528 | 2 | 0 | 0 |
| **NegCtrl1** | 1,406 | 55 | 3,097 | 59 | 2,486 | 12 | 1,340 | 7 |
| **NegCtrl2** | 256 | 21 | 2,511 | 23 | 1,672 | 3 | 1,229 | 2 |
| **NegCtrl3** | 74 | 8 | 154 | 16 | 103 | 4 | 68 | 4 |
| **NegCtrl4** | 137 | 4 | 142 | 5 | 111 | 1 | 0 | 0 |
| **NegCtrl5** | 12,145 | 130 | 13,493 | 304 | 10,466 | 20 | 0 | 0 |
| **ZymoStandCtrl1** | 3,169 | 4 | 6,033 | 77 | 4,401 | 3 | 3,024 | 2 |
| **ZymoStandCtrl2** | 10,469 | 28 | 19,132 | 178 | 13,771 | 5 | 9,590 | 7 |
| **ZymoStandCtrl3** | 3,675 | 4 | 6,722 | 74 | 5,010 | 3 | 3,480 | 2 |
| **ZymoStandCtr4** | 2,069 | 7 | 3,808 | 49 | 2,793 | 3 | 1,917 | 2 |
| **ZymoStandCtrl5** | 7,966 | 7 | 14,823 | 133 | 10,840 | 4 | 7,853 | 2 |
| **Total** | **9,926,235** | **27,920** | **10,108,666** | **66,381** | **8,060,929** | **1972** | **4,590,872** | **555** |
| **Mean** | 46,919 | 198 | 47,799 | 414 | 38,342 | 10 | 21,857 | 3 |
| **Median** | 15,076 | 125 | 15,514 | 143 | 11,357 | 8 | 4,435 | 3 |
| **Min** | 0 | 0 | 4 | 3 | 0 | 0 | 0 | 0 |
| **Max** | 4,528,706 | 5,185 | 4,581,763 | 27,218 | 3,788,697 | 62 | 2,474,314 | 19 |
| **Std.** | 342,297 | 458 | 346,526 | 2,125 | 285,514 | 8 | 185,135 | 3 |
