## Supplementary material for "Dataset complexity impacts both MOTU delimitation and biodiversity estimates in eukaryotic 18S rRNA metabarcoding studies": Table S1

| **Samples** | **UCLUST** | | **VSearch** | | **DADA2** | | **Deblur** | |
| --- | --- | --- | --- | --- | --- | --- | --- | --- |
|  | **Reads** | **OTUs** | **Reads** | **OTUs** | **Reads** | **ASVs** | **Reads** | **ASVs** |
| **A1.02.R1** | 111,295 | 3,946 | 111,947 | 3,763 | 94,021 | 256 | 45,393 | 304 |
| **A1.04.R1** | 146,089 | 4,288 | 147,357 | 5,594 | 120,123 | 148 | 52,823 | 176 |
| **A1.06.R1** | 49,695 | 3,125 | 55,733 | 3,553 | 44,006 | 224 | 22,973 | 271 |
| **A1.06.R2** | 139,795 | 4,505 | 140,860 | 5,700 | 122,454 | 347 | 59,027 | 357 |
| **A1.100** | 6,772 | 291 | 10,640 | 197 | 7,350 | 11 | 3,978 | 11 |
| **A1.20** | 75,253 | 2,792 | 84,402 | 2,911 | 72,297 | 232 | 36,864 | 271 |
| **A1.200** | 54,754 | 2,342 | 55,412 | 2,666 | 48,491 | 137 | 21,961 | 167 |
| **A1.350** | 151,520 | 2,931 | 152,204 | 4,647 | 121,171 | 54 | 62,554 | 74 |
| **A1.50** | 75,051 | 2,849 | 61,718 | 2,396 | 35,394 | 26 | 21,081 | 42 |
| **A2.100** | 141,759 | 4,100 | 136,241 | 3,303 | 113,966 | 110 | 57,435 | 134 |
| **A2.20** | 174,326 | 3,126 | 168,851 | 2,596 | 145,441 | 65 | 76,762 | 73 |
| **A2.200** | 75,165 | 2,533 | 78,061 | 3,121 | 67,138 | 138 | 33,517 | 153 |
| **A2.350** | 68,927 | 2,631 | 68,115 | 1,977 | 56,003 | 124 | 34,630 | 116 |
| **A2.50** | 155,463 | 3,278 | 140,299 | 4,265 | 79,856 | 32 | 45,889 | 40 |
| **A4.100** | 17,114 | 802 | 24,584 | 633 | 15,722 | 47 | 9,659 | 45 |
| **A4.20** | 69,898 | 1,874 | 69,241 | 1,930 | 52,551 | 105 | 32,824 | 126 |
| **A4.35** | 111,538 | 2,977 | 112,691 | 3,366 | 98,874 | 152 | 47,158 | 178 |
| **A4.50** | 64,613 | 3,076 | 62,905 | 1,983 | 48,895 | 86 | 23,530 | 101 |
| **A5.100** | 158,899 | 4,330 | 164,349 | 5,292 | 136,219 | 157 | 52,697 | 182 |
| **A5.20** | 82,457 | 4,445 | 79,274 | 3,839 | 63,708 | 159 | 30,317 | 213 |
| **A5.200** | 76,228 | 3,565 | 76,689 | 3,195 | 68,204 | 231 | 32,883 | 247 |
| **A5.35** | 228,262 | 3,656 | 219,323 | 5,499 | 151,136 | 101 | 82,122 | 118 |
| **A5.350** | 25,824 | 1,278 | 26,225 | 990 | 22,539 | 119 | 12,097 | 103 |
| **A5.50** | 100,687 | 2,662 | 103,100 | 1,945 | 83,970 | 66 | 44,748 | 89 |
| **A6.100** | 74,611 | 4,221 | 73,133 | 3,146 | 57,404 | 218 | 26,665 | 268 |
| **A6.20** | 103,279 | 1,377 | 101,910 | 1,669 | 87,003 | 36 | 48,310 | 36 |
| **A6.37** | 79,748 | 3,891 | 79,935 | 3,599 | 70,512 | 228 | 35,870 | 278 |
| **AGK.01** | 90,719 | 3,708 | 92,994 | 3,523 | 77,705 | 195 | 32,018 | 249 |
| **AGK.02** | 49,544 | 2,405 | 49,873 | 1,624 | 42,312 | 109 | 18,919 | 148 |
| **ALX.01** | 141,183 | 3,962 | 140,677 | 4,941 | 114,831 | 130 | 53,679 | 187 |
| **AMN.01** | 22,326 | 1,049 | 74,854 | 801 | 68,501 | 84 | 39,557 | 98 |
| **B1.200A.R2** | 142,818 | 1,420 | 144,404 | 1,122 | 126,901 | 57 | 82,544 | 68 |
| **B1.200B.R2** | 135,940 | 3,666 | 136,858 | 2,849 | 123,299 | 170 | 65,692 | 179 |
| **B1.200C.R1** | 4,688 | 712 | 9,528 | 672 | 4,522 | 61 | 2,133 | 84 |
| **B1.200C.R2** | 72,457 | 3,227 | 72,089 | 2,264 | 63,237 | 159 | 37,766 | 197 |
| **B1.500A.R1** | 142,097 | 3,279 | 143,138 | 3,638 | 115,450 | 78 | 76,196 | 101 |
| **B1.500B.R1** | 90,815 | 2,755 | 91,905 | 2,199 | 76,243 | 137 | 34,401 | 164 |
| **B1.500B.R2** | 78,605 | 2,719 | 75,789 | 1,970 | 61,931 | 63 | 38,904 | 85 |
| **B2.50.R2** | 44,511 | 2,827 | 53,475 | 2,666 | 46,177 | 181 | 25,580 | 230 |
| **BF003** | 113,249 | 2,817 | 112,781 | 2,470 | 86,688 | 103 | 52,228 | 118 |
| **BF005** | 100,017 | 4,748 | 101,880 | 4,735 | 85,879 | 222 | 36,026 | 244 |
| **BF011.R2** | 165,435 | 3,591 | 176,176 | 3,532 | 158,160 | 362 | 91,946 | 352 |
| **BF015** | 77,300 | 3,647 | 78,438 | 3,172 | 68,089 | 184 | 30,300 | 225 |
| **BNK.01** | 92,686 | 2,602 | 95,020 | 1,912 | 84,574 | 179 | 43,809 | 198 |
| **BNK.02** | 64,500 | 3,078 | 67,912 | 2,110 | 61,913 | 438 | 24,783 | 361 |
| **BNK.03** | 74,795 | 2,580 | 76,804 | 2,459 | 66,220 | 165 | 30,193 | 185 |
| **BNK.04** | 28,447 | 2,098 | 67,365 | 1,893 | 58,437 | 230 | 30,464 | 233 |
| **BNK.05** | 70,723 | 3,365 | 77,164 | 2,782 | 69,523 | 544 | 25,745 | 433 |
| **BNK.06** | 94,656 | 2,324 | 94,340 | 2,969 | 81,643 | 128 | 41,968 | 149 |
| **BNK.07** | 82,393 | 3,974 | 85,421 | 4,526 | 71,477 | 266 | 29,179 | 257 |
| **BPT.01** | 81,497 | 2,203 | 82,778 | 1,810 | 73,167 | 112 | 42,622 | 132 |
| **BPT.02** | 122,480 | 3,626 | 118,009 | 3,870 | 101,513 | 204 | 53,690 | 282 |
| **BPT.03** | 134,371 | 3,222 | 129,154 | 2,613 | 110,148 | 159 | 55,029 | 230 |
| **BPT.04** | 66,023 | 3,789 | 67,835 | 3,333 | 60,290 | 294 | 27,167 | 391 |
| **BPT.05** | 100,330 | 4,713 | 101,826 | 4,163 | 77,185 | 267 | 36,098 | 298 |
| **CPY.02** | 166,236 | 6,136 | 160,444 | 6,878 | 132,985 | 254 | 63,972 | 302 |
| **CPY.03** | 148,295 | 3,887 | 147,675 | 4,894 | 118,870 | 202 | 50,652 | 214 |
| **DF002.R1** | 10,611 | 1,133 | 15,133 | 1,145 | 9,909 | 128 | 5,208 | 193 |
| **DF002.R2** | 233,685 | 4,309 | 251,131 | 4,213 | 222,960 | 451 | 136,997 | 434 |
| **DF006.R1** | 34,594 | 1,909 | 37,200 | 1,435 | 30,929 | 100 | 16,030 | 138 |
| **DF006.R2** | 106,800 | 2,863 | 109,738 | 2,697 | 83,471 | 113 | 44,150 | 157 |
| **DF007.R2** | 8,356 | 599 | 22,897 | 748 | 7,083 | 37 | 3,228 | 48 |
| **DOL.01** | 107,722 | 2,481 | 109,907 | 2,354 | 91,870 | 113 | 49,398 | 139 |
| **DOL.03** | 109,202 | 3,722 | 117,938 | 3,065 | 102,496 | 361 | 46,929 | 314 |
| **DOL.04** | 53,669 | 2,687 | 54,887 | 2,317 | 46,553 | 185 | 21,833 | 226 |
| **DOL.05** | 58,693 | 1,923 | 55,338 | 1,738 | 45,337 | 75 | 29,338 | 89 |
| **DWT.10.R1** | 32 | 19 | 1,094 | 117 | 30 | 1 | 8 | 2 |
| **DWT.10.R2** | 63,255 | 3,347 | 62,684 | 3,079 | 52,349 | 120 | 27,117 | 129 |
| **ESC.03** | 122,716 | 4,555 | 118,559 | 3,573 | 87,233 | 165 | 45,993 | 165 |
| **ESC.04** | 83,994 | 3,580 | 79,308 | 3,276 | 59,017 | 170 | 21,103 | 180 |
| **FRK.01** | 111,316 | 4,442 | 105,409 | 3,997 | 75,678 | 199 | 41,453 | 227 |
| **FRK.02** | 50,648 | 3,096 | 50,990 | 2,938 | 32,731 | 116 | 13,611 | 138 |
| **FRK.03** | 205,293 | 2,380 | 204,044 | 2,424 | 174,386 | 111 | 92,381 | 138 |
| **FRK.04** | 88,009 | 1,712 | 84,037 | 1,719 | 69,488 | 44 | 42,267 | 36 |
| **FRK.05** | 113,080 | 6,137 | 106,933 | 4,798 | 79,095 | 207 | 38,882 | 235 |
| **FRK.06** | 30,601 | 2,134 | 31,380 | 1,595 | 23,732 | 66 | 11,453 | 82 |
| **FRK.07** | 70,960 | 3,455 | 69,704 | 3,003 | 52,334 | 111 | 25,932 | 144 |
| **GRY.02.R1** | 69,292 | 2,969 | 70,343 | 2,425 | 62,526 | 150 | 29,166 | 201 |
| **GRY.02.R2** | 49,137 | 2,254 | 50,071 | 1,735 | 43,414 | 114 | 26,615 | 154 |
| **GRY.04.R1** | 103,992 | 3,490 | 106,170 | 4,978 | 88,677 | 126 | 34,989 | 163 |
| **GRY.04.R2** | 49,514 | 3,452 | 48,638 | 3,524 | 39,335 | 108 | 16,520 | 139 |
| **GRY.06.R1** | 80,940 | 2,597 | 82,424 | 2,732 | 67,416 | 165 | 41,062 | 222 |
| **GRY.06.R2** | 244,357 | 4,083 | 251,495 | 4,545 | 223,875 | 575 | 132,574 | 498 |
| **HC010.R2** | 110,787 | 2,459 | 118,812 | 2,731 | 66,217 | 66 | 28,977 | 83 |
| **HC020** | 19,165 | 903 | 18,716 | 551 | 15,970 | 25 | 9,041 | 31 |
| **HC028** | 71,561 | 1,518 | 72,846 | 1,679 | 62,004 | 25 | 31,209 | 29 |
| **HC030** | 92,859 | 3,634 | 90,921 | 3,676 | 70,066 | 107 | 38,178 | 110 |
| **HN015.R2** | 53 | 35 | 1,396 | 149 | 90 | 3 | 9 | 4 |
| **HN016** | 108,101 | 3,412 | 112,932 | 4,494 | 74,662 | 91 | 33,378 | 96 |
| **HN025** | 89,423 | 3,275 | 90,225 | 2,828 | 63,362 | 96 | 31,312 | 112 |
| **HN029.R2** | 128,173 | 1,713 | 131,559 | 2,361 | 119,311 | 60 | 62,439 | 67 |
| **HS001** | 82,819 | 1,948 | 81,757 | 1,463 | 68,868 | 53 | 34,975 | 54 |
| **HS009** | 92,445 | 3,589 | 94,452 | 3,035 | 57,593 | 116 | 27,065 | 144 |
| **HS011** | 112,983 | 4,490 | 117,380 | 4,704 | 68,304 | 140 | 28,329 | 154 |
| **KF003.R2** | 44,801 | 1,852 | 47,230 | 1,556 | 39,291 | 100 | 21,770 | 124 |
| **KF011.R1** | 102,587 | 3,810 | 110,630 | 3,674 | 90,830 | 144 | 39,533 | 187 |
| **KF011.R2** | 128,531 | 3,318 | 138,442 | 3,522 | 82,210 | 74 | 41,406 | 90 |
| **KF015.R2** | 3,514 | 371 | 9,862 | 402 | 5,005 | 35 | 1,693 | 41 |
| **KF023.R1** | 140,535 | 4,540 | 147,146 | 4,106 | 124,114 | 245 | 51,958 | 303 |
| **KF023.R2** | 141,049 | 4,335 | 147,179 | 3,883 | 127,119 | 270 | 71,371 | 320 |
| **KUG.01** | 166,395 | 3,858 | 175,724 | 3,608 | 153,100 | 491 | 94,700 | 462 |
| **KUG.02** | 152,706 | 4,679 | 155,770 | 4,352 | 131,986 | 551 | 72,929 | 456 |
| **KUG.03** | 109,244 | 4,311 | 112,140 | 4,934 | 97,564 | 279 | 44,753 | 346 |
| **KUG.04** | 150,563 | 4,445 | 150,208 | 4,360 | 128,193 | 267 | 76,106 | 328 |
| **KUG.05** | 140,510 | 5,175 | 141,798 | 5,666 | 114,486 | 480 | 57,255 | 441 |
| **MNT.03** | 17,865 | 994 | 62,431 | 1,197 | 52,026 | 54 | 35,736 | 66 |
| **MNT.04** | 85,649 | 2,947 | 84,632 | 2,965 | 75,753 | 243 | 40,734 | 321 |
| **MNT.05** | 105,875 | 2,488 | 105,638 | 2,023 | 89,425 | 242 | 60,098 | 276 |
| **NTC.01** | 140,157 | 4,480 | 138,123 | 4,312 | 111,032 | 389 | 67,647 | 364 |
| **PWS.01** | 5,296 | 786 | 12,540 | 959 | 3,837 | 89 | 1,963 | 126 |
| **SF007.R1** | 170,971 | 4,112 | 173,760 | 4,071 | 151,623 | 313 | 89,353 | 329 |
| **SF007.R2** | 2,902 | 518 | 4,619 | 473 | 2,619 | 34 | 1,005 | 40 |
| **SF009.R1** | 116,578 | 4,751 | 117,359 | 4,119 | 93,072 | 225 | 53,431 | 269 |
| **SF009.R2** | 162,073 | 4,002 | 163,399 | 3,669 | 145,784 | 301 | 83,695 | 342 |
| **SF020** | 469 | 199 | 3,485 | 307 | 449 | 15 | 127 | 22 |
| **SMO.01** | 94,245 | 5,491 | 91,963 | 4,655 | 78,168 | 299 | 37,855 | 339 |
| **TBS.04.R1** | 147,468 | 4,387 | 143,980 | 4,244 | 118,163 | 193 | 70,144 | 226 |
| **TBS.04.R2** | 144,114 | 3,608 | 144,966 | 5,185 | 124,415 | 201 | 71,720 | 242 |
| **TBS.100** | 106,654 | 5,202 | 108,691 | 4,362 | 97,223 | 503 | 45,456 | 513 |
| **TBS.200** | 152,971 | 3,982 | 153,730 | 3,393 | 128,719 | 179 | 80,947 | 227 |
| **TBS.350** | 237,678 | 2,234 | 240,850 | 2,522 | 214,686 | 176 | 116,013 | 209 |
| **TBS.50** | 89,884 | 4,178 | 90,483 | 3,721 | 81,696 | 342 | 33,595 | 438 |
| **TF003.R1** | 148,160 | 4,245 | 149,112 | 3,719 | 127,082 | 255 | 75,321 | 322 |
| **WIS.01** | 89,587 | 4,374 | 87,927 | 3,888 | 72,459 | 227 | 40,093 | 275 |
| **WLK.01** | 757 | 181 | 4,543 | 339 | 714 | 21 | 270 | 23 |
| **WLK.02** | 15,104 | 1,648 | 15,349 | 1,351 | 13,154 | 71 | 4,353 | 81 |
| **WLK.03** | 201 | 78 | 2,997 | 219 | 294 | 8 | 66 | 13 |
| **Total** | **11,902,088** | **91,853** | **12,196,959** | **114,957** | **9,986,980** | **5,409** | **5,229,272** | **2,977** |
| **Mean** | 93,717 | 3,015 | 96,039 | 2,927 | 78,638 | 170 | 41,175 | 191 |
| **Median** | 90,815 | 3,222 | 91,905 | 2,969 | 73,167 | 140 | 37,766 | 176 |
| **Min** | 32 | 19 | 1,094 | 117 | 30 | 1 | 8 | 2 |
| **Max** | 244,357 | 6,137 | 251,495 | 6,878 | 223,875 | 575 | 136,997 | 513 |
| **Std.** | 53,935 | 1,344 | 52,742 | 1,433 | 45,634 | 123 | 26,350 | 119 |
